# Accelerated brain change in healthy adults is associated with genetic risk for Alzheimer’s disease and uncovers adult lifespan memory decline

**DOI:** 10.1101/2023.10.09.559446

**Authors:** James M. Roe, Didac Vidal-Piñeiro, Øystein Sørensen, Håkon Grydeland, Esten H. Leonardsen, Olena Iakunchykova, Mengyu Pan, Athanasia Mowinckel, Marie Strømstad, Laura Nawijn, Yuri Milaneschi, Micael Andersson, Sara Pudas, Anne Cecilie Sjøli Bråthen, Jonas Kransberg, Emilie Sogn Falch, Knut Øverbye, Rogier A. Kievit, Klaus P. Ebmeier, Ulman Lindenberger, Paolo Ghisletta, Naiara Demnitz, Carl-Johan Boraxbekk, the Alzheimer‘s Disease Neuroimaging Initiative, the Australian Imaging Biomarkers and Lifestyle Flagship Study of Ageing, Brenda Penninx, Lars Bertram, Lars Nyberg, Kristine B. Walhovd, Anders M. Fjell, Yunpeng Wang

**Affiliations:** Center for Lifespan Changes in Brain and Cognition (LCBC), Department of Psychology, University of Oslo, Norway; Norwegian Centre for Mental Disorders Research (NORMENT), Oslo University Hospital & Institute of Clinical Medicine, University of Oslo, Oslo, Norway; Department of Clinical Sciences Malmö, Lund University, Malmö, Sweden; Amsterdam UMC location Vrije Universiteit Amsterdam, Department of Psychiatry and Amsterdam Neuroscience, Amsterdam, The Netherlands; Department of Integrative Medical Biology, Umeå University, Sweden; Umeå center for Functional Brain Imaging, Umeå University, Sweden; Cognitive Neuroscience Department, Donders Institute for Brain, Cognition and Behavior, Radboud University Medical Center, Nijmegen, The Netherlands; Department of Psychiatry and Wellcome Centre for Integrative Neuroimaging, University of Oxford, Warneford Hospital, Oxford, United Kingdom; Center for Lifespan Psychology, Max Planck Institute for Human Development, Berlin, Germany; Max Planck UCL Centre for Computational Psychiatry and Ageing Research, Berlin, Germany, and London; Faculty of Psychology and Educational Sciences, University of Geneva, Switzerland; Faculty of Psychology, UniDistance Suisse, Brig, Switzerland; Danish Research Centre for Magnetic Resonance, Centre for Functional and Diagnostic Imaging and Research, Copenhagen University Hospital – Amager and Hvidovre, Copenhagen, Denmark; Institute for Clinical Medicine, Faculty of Medical and Health Sciences, University of Copenhagen, Copenhagen, Denmark. (CJ); Department of Radiation Sciences, Diagnostic Radiology, and Umeå Center for Functional Brain Imaging (UFBI), Umeå University, Umeå Sweden; Institute of Sports Medicine Copenhagen (ISMC) and Department of Neurology, Copenhagen University Hospital Bispebjerg, Copenhagen, Denmark; Lübeck Interdisciplinary Platform for Genome Analytics (LIGA), University of Lübeck, Germany; Computation Radiology and Artificial Intelligence, Department of Radiology and Nuclear Medicine, Oslo University Hospital, Oslo, Norway

**Keywords:** Lifespan neuroscience, ageing, longitudinal, polygenic risk, cognitive decline

## Abstract

Across healthy adult life our brains undergo gradual structural change in a pattern of atrophy that resembles accelerated brain changes in Alzheimer’s disease (AD). Here, using four polygenic risk scores for AD (PRS-AD) in a longitudinal adult lifespan sample aged 30 to 89 years (2-7 timepoints), we show that healthy individuals who lose brain volume faster than expected for their age, have a higher genetic AD risk. We first demonstrate PRS-AD associations with change in early Braak regions, namely hippocampus, entorhinal cortex, and amygdala, and find evidence these extend beyond that predicted by *APOE* genotype. Next, following the hypothesis that brain changes in ageing and AD are largely shared, we performed machine learning classification on brain change trajectories conditional on age in longitudinal AD patient-control data, to obtain a list of AD-accelerated features and model change in these in adult lifespan data. We found PRS-AD was associated with a multivariate marker of accelerated change in many of these features in healthy adults, and that most individuals above ∼50 years of age are on an accelerated change trajectory in AD-accelerated brain regions. Finally, high PRS-AD individuals also high on a multivariate marker of change showed more adult lifespan memory decline, compared to high PRS-AD individuals with less brain change. Our results support a dimensional account linking normal brain ageing with AD, suggesting AD risk genes speed up the shared pattern of ageing- and AD-related neurodegeneration that starts early, occurs along a continuum, and tracks memory change in healthy adults.

## Introduction

Advanced age is the primary risk factor for Alzheimer’s disease (AD) – the leading cause of dementia. Across healthy adult life and ageing, our brains undergo gradual and widespread structural changes^1–4^. Many of these changes are qualitatively similar to atrophy patterns seen in AD, suggesting shared vulnerability of brain systems in ageing and AD^5–7^. For example, medial temporal lobe regions including hippocampus and entorhinal cortex are amongst the earliest affected regions in AD in terms of structural atrophy and tau deposition^8,9^, and each exhibits accelerated structural loss from around ∼50 years of age. Prior to this, many brain structures exhibit slow but steady average volume reduction from early adulthood ^2,10–13^. Beyond specific regions, whole-cortical atrophy patterns are also largely shared between ageing and AD^4,6^, with characteristic temporo-parietal atrophy patterns in AD also found to a lesser degree in healthy people, including those at low AD risk^5,14^. It has been argued this parallel pattern is critical to understand^4^, because reported AD incidence increases exponentially after 65 years of age ^15,16^.

If brain regions vulnerable in AD also exhibit gradual change during adult life, healthy individuals at higher AD risk may show faster atrophy over extended age spans. Polygenic risk scores for AD (PRS-AD) calculated from AD risk variants found in genome-wide association studies (GWAS) provide a marker to test this; in AD patients, higher genetic AD risk links with longitudinal outcomes including faster brain and cognitive decline, earlier AD onset, and clinical progression^17–19^. In healthy adults, however, attempts to link genetic AD risk to alterations in brain structure have typically been cross-sectional and yielded mixed results^20–26^. For example, although many studies report no effect of *APOE-ε4* on cross-sectional hippocampal volumes^20–26^, recent large-scale studies found smaller hippocampal volumes in older *ε4* carriers^27,28^. However, evidence suggests smaller hippocampal volume as a function of genetic AD risk is evident in neonates^29^, children^2,30^ and young adults^31,32^, and longitudinal work suggests the effect of AD risk genes upon lower hippocampal volume is roughly equivalent from childhood to old age^2^. Further, many 70-year-olds have similar size brain measures to many 30-year-olds, and individual differences in brain structure at any age typically exceed the magnitude of change effects through ageing^2,33^. Hence, brain differences observed in older at-risk individuals may be ascribable to preexisting differences from early life. Consequently, only longitudinal designs are suited to examine whether elevated genetic AD risk confers a direct genetic effect on the slope of brain ageing across the healthy adult lifespan.

Longitudinal studies attempting to link brain changes to genetic AD risk in healthy adults have been inconclusive, often restricted to small samples of older adults^34–36^, and lifespan samples of healthy individuals with extensive follow-up over large age-spans are lacking. Small studies have reported group-level effects^34,35^ or no effect of *APOE-ε4* upon hippocampal change in healthy older adults^37^. Another study found evidence that PRS-AD related to hippocampal and entorhinal thinning in an older sample enriched for *APOE-ε4* and memory concerns, though did not report polygenic effects beyond *APOE*^36^. Additionally, in a large sample of healthy older individuals, hippocampal change was found to be greater in *APOE-ε4* carriers (N=748)^38^. However, a recent GWAS^39^ (N=15,640) observed that an association between *APOE* and faster hippocampal and amygdala change in ageing disappeared when accounting for disease status (notably, the sample included many AD cases). Thus, the effect of *APOE* upon brain change in candidate AD regions was seemingly driven by disease-related processes and not detected in healthy brains^39^. Moreover, the trajectories of genetically high-risk versus low-risk groups provide little evidence that genetic AD risk affects the slope of brain decline across the adult lifespan^2,24,25^. Individualized estimates of the degree to which a healthy person’s brain is changing more or less than expected for their age may be better suited to answer whether genetic AD risk impacts the slope of brain ageing in healthy adults.

Regions with greater brain atrophy in AD are encompassed within the Braak staging scheme^8,40^. This describes the spatiotemporal sequence of tau deposition^9,41^ – from a cortical entorhinal epicentre (stage I) to hippocampus (stage II), amygdala and inferior temporal cortex (stage III), and later to the rest of cortex^8,40^. This “AD signature”^9^ is not specific to AD but also found to a lesser degree in normal ageing^5,6,42^. Beyond this core set of regions with seemingly shared vulnerability to the effects of ageing and AD, many other brain features exhibit accelerated change in AD. Applying a data-driven approach to first delineate these in AD patients – combined with multivariate analyses using individualized brain change estimates in healthy adult lifespan data – may reveal new insights into whether genetic AD risk influences the slope of brain ageing in a select few or across many AD-relevant features in healthy adults.

Finally, several studies suggest that genetic AD risk is subtly related to longitudinal memory decline in healthy older adults^43–45^, and one adult lifespan study reported genetic AD risk was weakly associated with decline in a composite cognitive and memory score^46^. Thus, AD risk genes may influence differences in memory decline trajectories that are protracted through life and begin in early adulthood^45–48^. However, the extent to which AD risk genes influence brain and cognitive outcomes probably differs also between individuals at comparatively high genetic risk, which may explain why genetic risk alone is not highly predictive of cognitive change^46,49^. Given that individualized approaches to risk assessment are predicated on assessing the conjunction of risks, considering known genetic AD risk together with a brain risk marker may improve identification of individuals at higher AD risk, also in healthy adult lifespan data.

Here, in a healthy adult lifespan sample with frequent longitudinal follow-up, we establish that individuals changing more than their age would predict in AD-accelerated brain regions are at significantly enhanced genetic AD risk (2-7 timepoints, 1430 scans from 420 individuals aged 30 to 89 years). Using genome-wide significant single nucleotide polymorphisms (SNPs; *p* < 5×10^−8^) from four AD GWAS, we first 1) show that PRS-AD significantly associates with more age-relative change in early Braak stage regions. Next, to empirically identify brain features with accelerated change in AD, we run machine learning (ML) binary classification on the individual-specific slopes derived from longitudinal AD patient-control data from the Alzheimer’s Disease Neuroimaging Initiative (ADNI; scans = 4410, N = 978, 2-9 timepoints). Modelling change in these in our healthy adult lifespan sample, we 2) show that PRS-AD is significantly associated with change in many AD-accelerated brain features in healthy adults. In an independent replication sample with notably less follow-up (2-3 timepoints), we corroborate some of the observed PRS-AD associations with brain change in healthy adults. Last, we 3) show that high PRS-AD individuals also high on a multivariate brain change marker show greater drop-off in memory over the adult lifespan, compared to high PRS-AD individuals with less brain change. Thus, the conjunction of a multivariate brain change marker and known genetic risk helped identify a subset of individuals showing more memory decline over their healthy adult life (30-89 years).

## Results

### Age-relative brain change across the healthy adult lifespan associates with genetic AD risk *Univariate analyses: A priori ROI’s:*

To estimate age-relative brain change in adult lifespan data, we used all longitudinal scans fitting age-range and inclusion criteria (≥30 years of age; Methods). This allowed us to obtain the best-fitting age trajectory models from which we could subsequently estimate how much an individual’s change trajectory deviated from the population-average (i.e., from the level of change predicted given age), via individual-specific random slopes in a Generalized Additive Mixed Model of age (Methods). We first explored brain change in initial hippocampal ROI’s – Braak Stage II^50^. Fig. 1A-C shows the longitudinal lifespan trajectory, and individual-specific degree of absolute and age-relative change for the left hippocampus (see SI Fig. 1 for right hippocampus). As expected^2^, most individuals aged ≥30 years exhibited hippocampal volume loss, but to differing degrees, and very few individual-specific slopes were estimated to show growth over time. As also expected, the degree of absolute hippocampal change accelerated on average between the ages of 50 and 60 years. The degree of age-relative change was significantly associated with PRS-AD in the hypothesized negative direction: on average across the adult lifespan (30-89 years), individuals exhibiting more hippocampal loss than expected given their age had significantly higher PRS-AD. This genetic association was probed separately for the bilateral hippocampi (left: β = -.22, t(212) = -3.3, *p*=.001; right: β = -.16, t = 2.4, *p*=.015; [PRS-AD Jansen]; covariates: mean age, sex, N timepoints, and interval between first and last timepoint), and was significant using all four GWAS-derived scores. To ensure we were capturing ageing-specific effects at some point (see SI Fig. 1), we tested the association using change rates extracted from progressively older age-ranges (i.e., progressively discarding data from comparatively younger individuals; Methods). This also ensured that the analysis outcome was not based on a single arbitrary decision such as the age range to test the average association across^51,52^. FDR-correction was applied across all 576 PRS-AD tests reported in this analysis. We then tested whether surviving associations remained statistically significant at p<.05 using polygenic scores computed without *APOE* (PRS-AD^noAPOE^), assuming a 5% chance false positive rate per structure. Despite the progressively smaller sample size, all tested PRS-AD associations with age-relative hippocampal change (left and right) were significant at p<.05 [uncorrected] using all four scores (coloured points in Fig. 1E-F denote associations at p<.05 [uncorrected]; see lower panels for respective effect sizes). 31 of the 36 tests (86%) with age-relative left hippocampal change, and 25/36 (69%) with age-relative right hippocampal change, survived FDR-correction (see lower panels in Fig 1E-F; partial r^2^ effect size is shown for associations surviving FDR-correction). Using PRS-AD to predict absolute hippocampal change instead in comparable statistical models (i.e., also correcting for mean age), PRS-AD associations were also mostly significant after FDR correction (47/72 [65%] survived correction). Probing whether FDR-corrected associations with change remained after discounting the effect of *APOE* per each structure tested, 19/58 (33%) PRS-AD^noAPOE^ associations with left hippocampal change (age-relative or absolute) remained significant at p<.05, surpassing the 5% false positive rate expected by chance (black crosses in Fig. 1E-F denote partial r^2^ of PRS-AD^noAPOE^ where significant [*p*<.05]). For right hippocampus, 6/45 (13%) of the FDR-corrected associations remained significant at p<.05 with PRS-AD-^noAPOE^, also surpassing the chance false positive rate (Fig. 1F). Post-hoc tests confirmed the impression that the estimated regression coefficients became more negative as the age subset steadily comprised only older individuals (Fig. 1E-F); on average across change metrics, each increasing age subset was associated with a reduction in the negative beta coefficient of -.026 for left hippocampus (t = -14.1; *p*_*perm*_ = 9.9e^-4^), and -.023 for right hippocampus (t = -15.4; *p*_*perm*_ = 9.9e^-4^). Alternative post-hoc analyses dependent on power across the full age-range (30-89 years) found significant PRS-AD × age (mean) interactions upon age-relative change in left and right hippocampi for all four scores but these did not survive multiple comparison correction (SI Table 1; SI Fig. 3).

**FIGURE 1.**
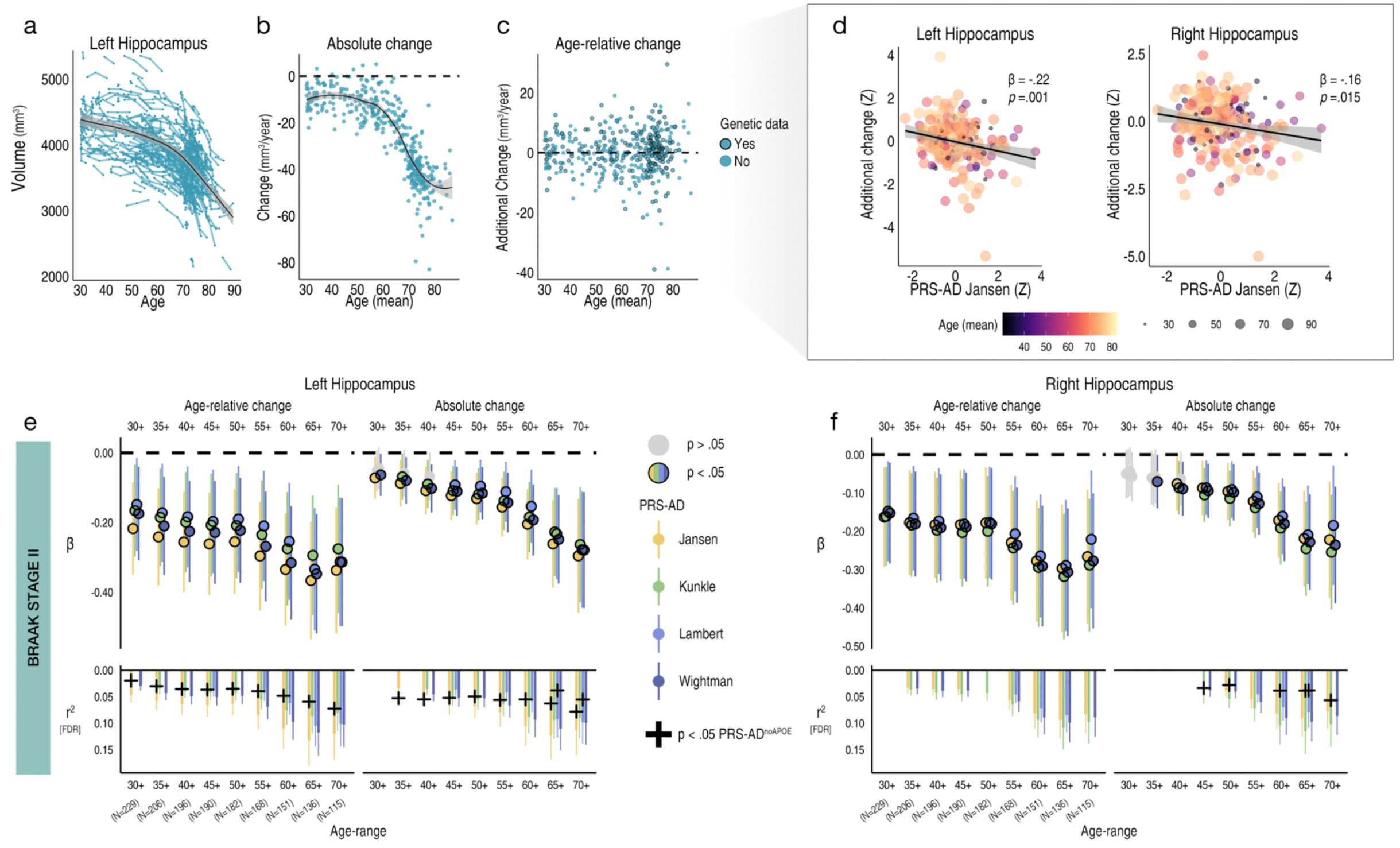
Hippocampal change in healthy adults associates with genetic AD risk. Exclusively longitudinal data was used to estimate individual-specific age-relative and absolute change in hippocampus (Braak stage II), modelling the adult lifespan trajectories using GAMMs with random slopes. **a** Adult lifespan trajectory for left hippocampus from 30-89 years (data corrected for sex and scanner). Lines connect longitudinal observations per participant. **b** Absolute change per individual (datapoints) in left hippocampus as a function of their mean age across timepoints. **c** Estimated age-relative change per individual in left hippocampus (i.e., individual-specific slopes) as a function of their mean age across timepoints. Black stroke indicates whether or not genetic data was available per participant and thus whether the data-point was included in the PRS-AD association tests. **d** More negative age-relative change in both left and right hippocampus was associated with significantly higher PRS-AD on average across the full adult lifespan sample with genetic data (30-89 years; N=229; association visualized for one score [Jansen]); age and covariate-corrected [Methods]; colour and datapoint size depicts mean age). **e-f** PRS-AD associations with age-relative change (left facet) and absolute change (right facet) in left (E) and right (F) hippocampus, using the four GWAS-derived scores, tested for progressively older age-ranges to ensure capture of ageing-specific effects (i.e., moving from left to right on the X-axis, the leftmost age-range represents the association tests across the full adult lifespan on average [30-89 years; N=229], whereas the right-most age-range shows the associations tested in only the oldest adults [70-89 years]; standardized β). Significant associations at *p* < .05 are depicted in colour (upper panels). For associations surviving FDR-correction, partial r^2^ of PRS-AD is shown (lower panels). Where the association survived FDR-correction, we retested the association after removing *APOE* (PRS-AD^noAPOE^). Partial r^2^ of PRS-AD^noAPOE^ is depicted by a black cross if the FDR-corrected association remained significant (*p* < .05). Ribbons and error bars depict 95% CI.

We then repeated the procedure for Braak stage I (entorhinal) and III regions (subcortical and cortical ROIs; Methods). For Braak stage I, we observed no significant PRS-AD associations with change (age-relative or absolute) in left entorhinal cortex, but observed several significant associations with each in right entorhinal cortex, 5 of which survived correction (Fig. 2A). 3 of these FDR-corrected associations remained significant at p<.05 with PRS-AD^noAPOE^ (using absolute change; lower panels in Fig. 2A), surpassing the false positive rate. Post-hoc tests confirmed that the estimated regression coefficients became more negative as the age subset comprised only older individuals for right (beta reduction = -.013, t = -7.8, *p*_*perm*_ = .018) but not for left entorhinal cortex (beta reduction = -.008, t = -5.0, *p*_*perm*_ = .10). However, alternative post-hoc analyses across the full age-range (30-89 years) found no PRS-AD × age (mean) interactions upon age-relative entorhinal change, suggesting our data may have been underpowered to detect a two-way continuous interaction (SI Fig. 3; SI Table 1).

**FIGURE 2.**
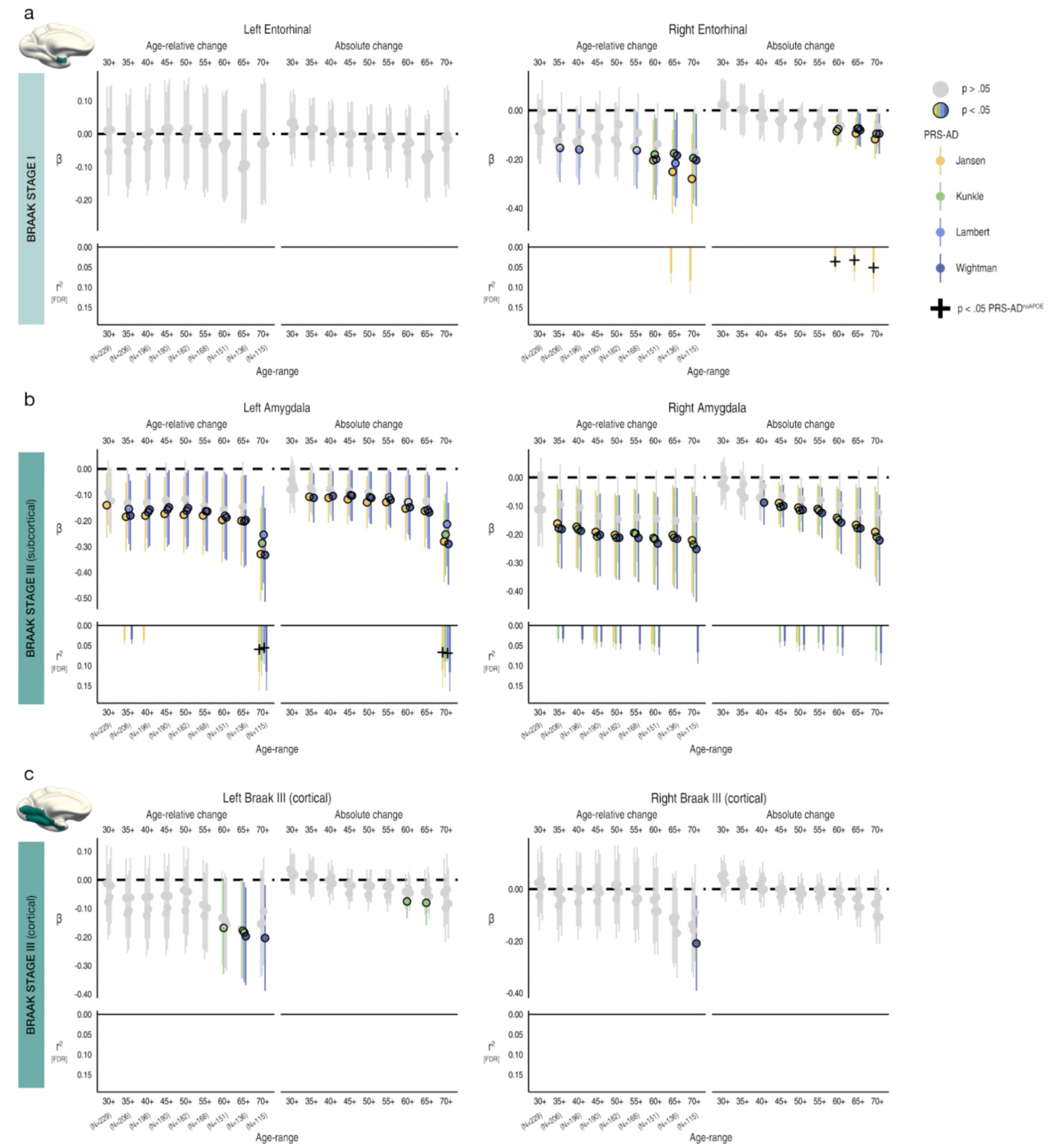
Change in early Braak stage regions in healthy adults associates with genetic AD risk. PRS-AD associations with age-relative change and absolute change in brain regions encompassed within **a** Braak stage I (entorhinal) and **b-c** Braak stage III regions (amygdala and inferior temporal cortical ROI), using the four GWAS-derived scores, tested for progressively older age-ranges to ensure capture of ageing-specific effects (i.e., moving from left to right on the X-axis, the leftmost age-range represents the association across the full adult lifespan on average [30-89 years; N=229], whereas the rightmost age-range shows the associations tested in only the oldest adults [70-89 years]; standardized β). Significant associations at *p* < .05 are depicted in colour (upper panels). Partial r^2^ of PRS-AD is shown for all associations surviving FDR-correction (lower panels; lower panels in A [left] and C [left and right] are correctly empty because no association survived correction). Where the association survived FDR-correction, we retested the association after removing *APOE* (PRS-AD^noAPOE^). Partial r^2^ of PRS-AD^noAPOE^ is depicted by a black cross if the FDR-corrected association remained significant (*p* < .05). Error bars depict 95% CI.

For the subcortical Braak Stage III region (amygdala), we similarly observed negative associations between age-relative change in left and right amygdala and PRS-AD (Fig. 2B), 21 of which were significant after FDR correction, and using absolute change instead yielded similar results (15 surviving associations). 4/11 (36%) PRS-AD^noAPOE^ associations remained significant at p<.05 for left amygdala (surpassing the false positive rate), whereas no associations with right amygdala change remained after excluding *APOE*. The estimated regression coefficients became stronger as the age subset comprised only older individuals (beta reduction left amygdala = -.018, t = -7.6, *p*_*perm*_ = .002; right amygdala = -.019, t = 10.0 *p*_*perm*_ = .004), though alternative analyses dependent on power across the full age-range found no post-corrected significant PRS-AD × age (mean) interactions upon age-relative change in amygdala (SI Fig. 3; SI Table 1). For the cortical component of Stage III, none of the tested PRS-AD associations with change in left or right cortex survived correction (Fig. 2C), the regression coefficients became stronger as the age subset comprised only older individuals in each (beta reduction left cortex = -.011, t = -6.2, *p*_*perm*_ = .013; right cortex = -.013, t = -6.3; *p*_*perm*_ = .004), and we found no significant PRS-AD × age (mean) interactions in alternative analyses across the full age-range (SI Fig. 3; SI Table 1).

### Multivariate analyses: data-driven features exhibiting accelerated change in AD

Given the univariate results, we expected that multivariate measures of change would be better suited to detect PRS-AD associations with brain change in healthy adults. Thus, we sought to empirically obtain a list of brain features with accelerated change in AD, then test whether multivariate change across these features relates to PRS-AD in the LCBC healthy adult lifespan discovery sample (Methods). First, in longitudinal AD patient-control data from ADNI (SI Table 2), we defined two longitudinal groups we could be maximally confident consisted of healthy individuals and those succumbing to AD based on diagnosis: *NC-long* consisted of normal controls consistently classed as healthy over time, whereas *AD-long* comprised all individuals with an AD diagnosis by their final timepoint (Fig. 3A; Methods). Then, in 364 features we modelled a GAMM of age (irrespective of group), and entered the individual-specific slopes into ML binary classification (Fig. 3B). Group differences in slopes (age-relative change) were in the expected direction (Fig. 3C). The top features deemed most important for separating *AD-long* from *NC-long* individuals based on age-relative change in ADNI included many well-known AD brain vulnerabilities (e.g., ventricles, medial temporal and temporo-parietal regions; see Fig. 4A; though our intention was not to refine prediction of AD cases, we note the model achieved an area under the curve [AUC] of .952 in independent data from the Australian Imaging Biomarker & Lifestyle Flagship Study of Ageing [AIBL]; Fig. 3D-F; SI Fig. 4).

**FIGURE 3.**
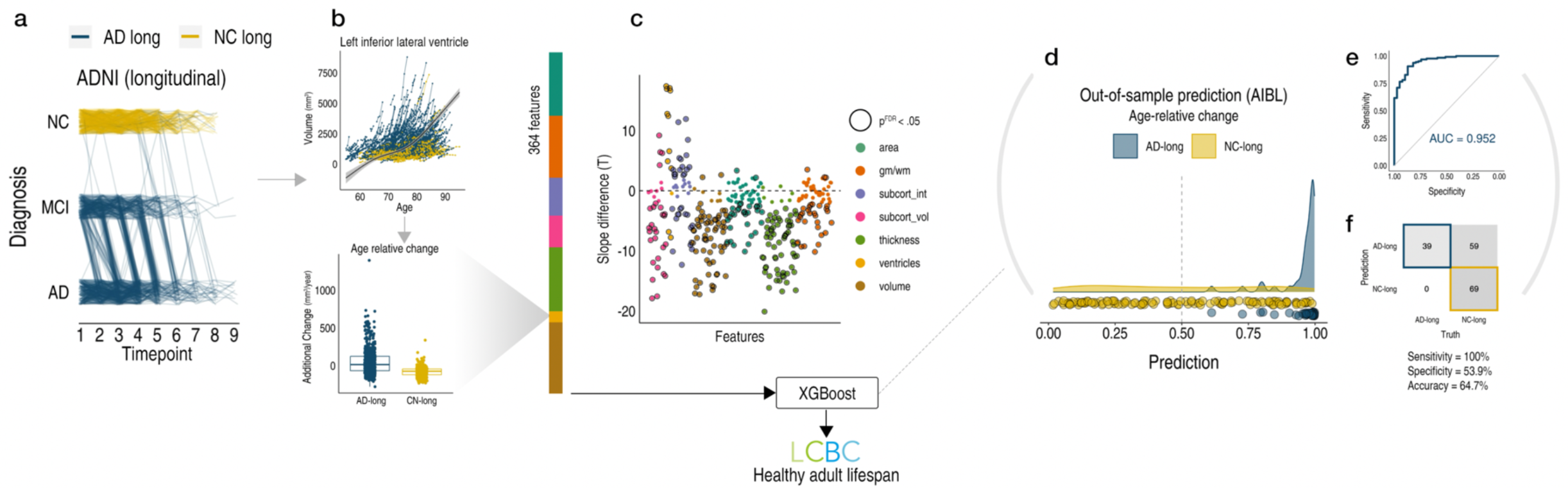
Visualization of longitudinal AD analysis pipeline. **a** Longitudinal grouping in ADNI data. X-axis denotes the scan observations across timepoints used in the final sample. Each line represents a participant and colour denotes longitudinal group membership. Single-timepoint ADNI diagnoses (Y-axis; NC normal controls, MCI mild cognitive impairment, AD Alzheimer’s disease) were used to define two longitudinal groups of AD and NC individuals (AD-long; N = 606, obs = 2730; NC-long, N = 372; obs = 1680). *NC-long* individuals were classified as healthy at every timepoint whereas *AD-long* individuals were diagnosed with AD by their final timepoint (Methods). Single-timepoint MCI diagnoses were considered only for the purpose of defining the longitudinal AD group. Note that because the grouping used all diagnosis observations (i.e., not only scan observations), trajectories of individuals that appear to end with a NC or MCI diagnosis nevertheless correspond to individuals with an AD diagnosis by their final timepoint, as do those seemingly reverting (Methods). **b** GAMMs of Age (across groups; upper plot) were used to model age-relative change (individual-specific slopes) in 364 brain features (shown for one example feature). The ADNI-derived individual-specific slopes were then used as input to machine learning binary classification using XGBoost ^53^ **c** Most features exhibited significant group-differences in age-relative change between *AD-long* and *NC-long* as expected (datapoints denote t-statistics for t-tests; black stroke indicates significant associations at p(FDR)<.05). **d-f** Out-of-sample prediction for the binary classifier (AIBL data; SI Fig. 4) including receiver operator curve (d), confusion matrix and performance metrics (e). The purpose of the classification procedure was to empirically derive brain features with accelerated change in AD, to use these in healthy adult lifespan data.

**FIGURE 4.**
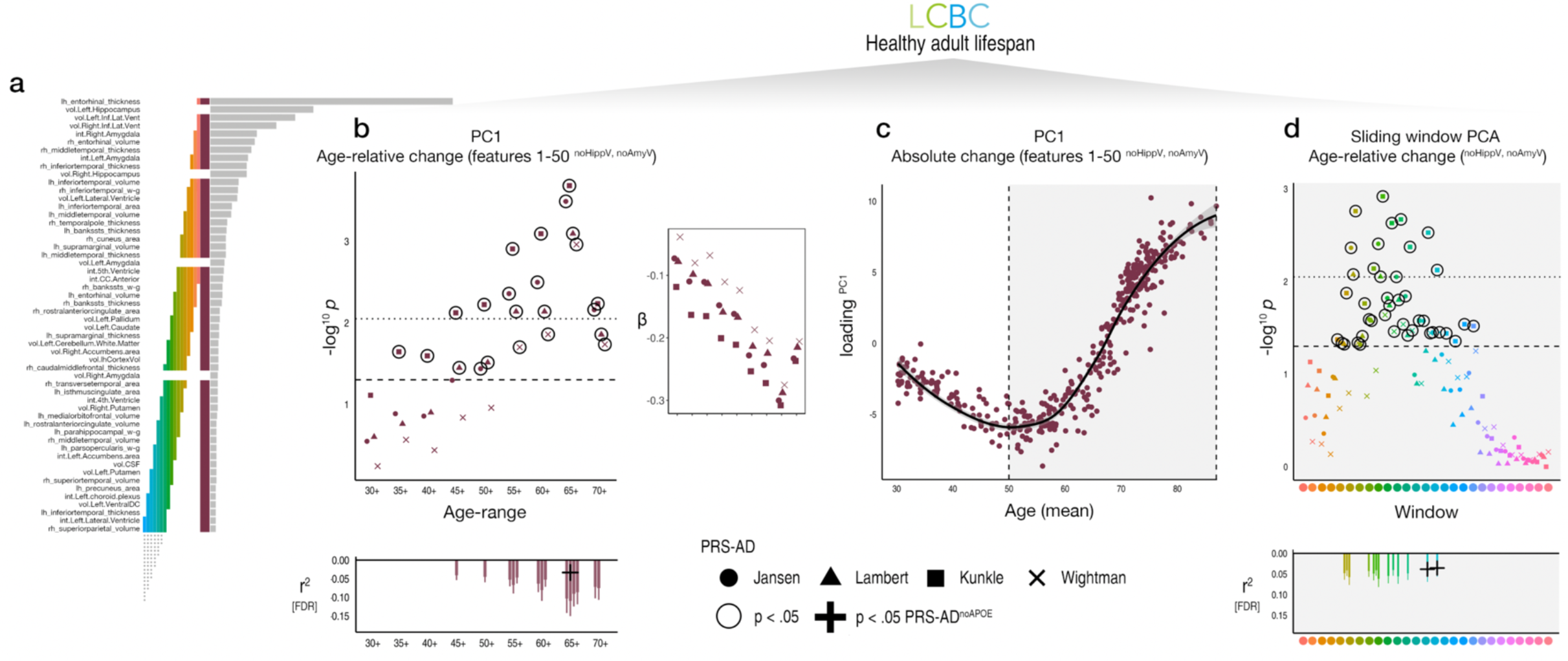
ADNI-derived features applied to the healthy adult lifespan. **a** Top features for classifying *AD-long* from *NC-long* individuals in ADNI data based on age-relative change. Coloured bars indicate feature selections across which we calculated PC1 and link with the subsequent plots. **b** PRS-AD associations in healthy adult lifespan data (LCBC sample) using the principal component of age-relative change across the top 50 brain features with accelerated change in AD (excluding hippocampal and amygdala volumes; PC1^relChange^; maroon bar in a). Datapoints show (-log10) p-values for the association with PC1^relChange^, tested at progressively older age-ranges, for all four scores. Dashed line indicates p = .05, and datapoints with black stroke denote significant PRS-AD associations at p<.05. Datapoints above the dotted line are significant at p(FDR)<.05. Smaller right inset plot shows the standardized Beta values as a function of age-range (Betas inversed to be negative due to the non-directional nature of PCA). Bottom plot shows partial r^2^ of PRS-AD for all associations surviving FDR-correction. Where the association survived FDR-correction, we retested the association after removing *APOE* (PRS-AD^noAPOE^). Partial r^2^ of PRS-AD^noAPOE^ is depicted by a black cross if the FDR-corrected association remained significant (*p* < .05). Error bars depict 95% CI. **c** The principal component of absolute change across the top 50 brain features with accelerated change in AD (excluding hippocampal and amygdala volume; maroon bar in a), plotted as a function of mean age across timepoints. Accelerated brain change in AD-accelerated features was evident between ages 50-60. Note that since the y-axis represents change, the slope of the curve represents acceleration (see also SI Figs. 10-11). **d** PCA-based sliding window analysis within the age-range 50-89 years. Colours and order correspond to the coloured bars in a, which show the selection of features across which the principal component of age-relative change was calculated and used to test associations with PRS-AD. Dashed line indicates p = .05, and datapoints with black stroke denote significant PRS-AD associations at p < .05. Datapoints above the dotted line are significant at p(FDR)<.05. lh=left hemisphere, rh=right hemisphere, vol=volume (subcortical); int=intensity (sub-cortical); w-g=grey/white matter contrast. Subcortical features (aseg) are delineated with “.”, whereas cortical features (aparc) are delineated with “_”.

In the LCBC healthy adult lifespan discovery sample, we then calculated the principal component of agerelative change across the first 50 features with model-implied importance (PC1^relChange^; hippocampal and amygdala volumes were not included in PC1^relChange^ to ensure these did not drive the multivariate effect; see the maroon bar in Fig. 4A; explaining 13% variance). As hypothesized, 14 of the 36 tested associations relating PC1^relChange^ to PRS-AD were FDR-corrected significant (Fig. 4B; FDR-correction applied across all 144 PRS-AD tests in this analysis). Again, post-hoc tests confirmed that the estimated regression coefficients became stronger as the age subset comprised only older individuals (beta reduction = -.023, t = -9.9, *p*_*perm*_ = .002) and alternative analyses across the full age-range found post-corrected significant PRS-AD × age (mean) interactions upon PC1^relChange^ using all four scores (SI Table 3). Next, to determine the age at which brain change in AD-accelerated features starts increasing in healthy adults, we took the principal component of absolute change across the same set of 50 features (PC1^absChange^; explaining 45% variance) plotted as a function of mean age (Fig. 4C). The results suggested that all individuals were on a trajectory of change in AD features that showed onset of accelerated change around age ∼50 in healthy adults (Fig. 4C; see SI Fig. 10 for derivative plots). Further, change trajectories were steepest in features most important for separating AD-patients from controls (SI Fig. 6). To ensure that the multivariate associations were not driven by one or a few brain features, we ran a sliding window PCA within the 50-89 year age-range (Methods). PRS-AD associations with age-relative change were evident when calculating PC1 across many combinations of features, including those relatively lower down in terms of model importance (coloured bars in Fig. 4A denote feature windows for the PCA and link with the coloured points denoting p-values for the PRS-AD associations in Fig. 4D; see SI Fig. 7 for correlations between features). 13 of the tested associations were significant after FDR correction, illustrating that multivariate change across many AD-accelerated features relates to PRS-AD in healthy adults (Fig. 4D). The data suggested that PRS-AD associations derived via this method were largely though not entirely driven by *APOE* (3 of the 27 [11%] FDR-corrected tests remained significant at p < .05 using PRS-AD^noAPOE^, surpassing the 5% false positive rate; lower panels in Fig. 4).

As a final proof-of-principle, we directly applied the ADNI-derived model weights to the LCBC healthy adult lifespan discovery sample. This prediction incorporates information from the weights of all 364 features (Methods). The dependent variable was the model-implied log odds of having AD (probAD^relChange^; Methods). Importantly, because the model was trained on an index of relative brain change conditional on age, the logistic prediction applied to the healthy adult lifespan data cannot be interpreted in terms of its implied binary outcome (i.e., AD/no-AD). This is because the model could assign the same probability of having AD to a hypothetical 30-year-old with an estimated additional brain loss of 10mm^3^/year as to a 60-year-old with the same additional brain loss, even though change and AD risk are higher in the 60-year-old, because change is over and above the mean brain loss anticipated at age 60 (see Fig. 1C). We nevertheless hypothesized the learned model weights would be useful, and would relate to PRS-AD in a similar way to the raw age-relative change values in specific features. As expected, almost all of the tested PRS-AD associations with probAD^relChange^ were significant at p<.05, 14 of which survived correction; see SI Fig. 5B). Repeating all steps of the model estimation procedure using absolute change instead (from hyperparameter estimation to prediction; AUC = .933 in unseen data from AIBL), we found far fewer significant PRS-AD associations with probAD^absChange^ (7 survived correction; SI Fig. 5C-D), suggesting relative change is a superior marker for capturing individual differences in brain ageing. Again, the data indicated PRS-AD associations derived by this method were largely though not entirely driven by *APOE* (8 [38%] of the FDR-corrected tests with change remained significant using PRS-AD^noAPOE^; SI Fig. 5; FDR-correction applied across all 72 PRS-AD tests in this analysis).

### Replication analysis

To reduce the number of tests, in an independent adult lifespan replication sample with fewer follow-up points (2-3 timepoints; Lifebrain replication sample), we tested PRS-AD associations using hippocampal and amygdala change, and the principal component of age-relative change across the first 50 AD-accelerated features, not including hippocampal or amygdala volume (i.e., PC1^relChange^; Fig. 4A). For hippocampus, we observed similarly negative effects, 22 of which were significant for age-relative change (p < .05 [uncorrected]; 31 for absolute change; Fig. 5A). Similar to the discovery sample, PRS-AD effects on age-relative hippocampal change were larger than absolute change, and often remained significant after discounting *APOE* (black crosses in Fig. 5A denote partial r^2^ for PRS-AD^noAPOE^ where this remained significant). For amygdala, we observed no significant PRS-AD associations within any age-range, and we also observed no significant associations with PC1^relChange^ (Fig. 5B-C). However, like the discovery sample, all healthy individuals lay on a trajectory of accelerated change in AD features, with a similar onset of acceleration around the age of 50 years (Fig. 5D; SI Fig. 10).

**FIGURE 5.**
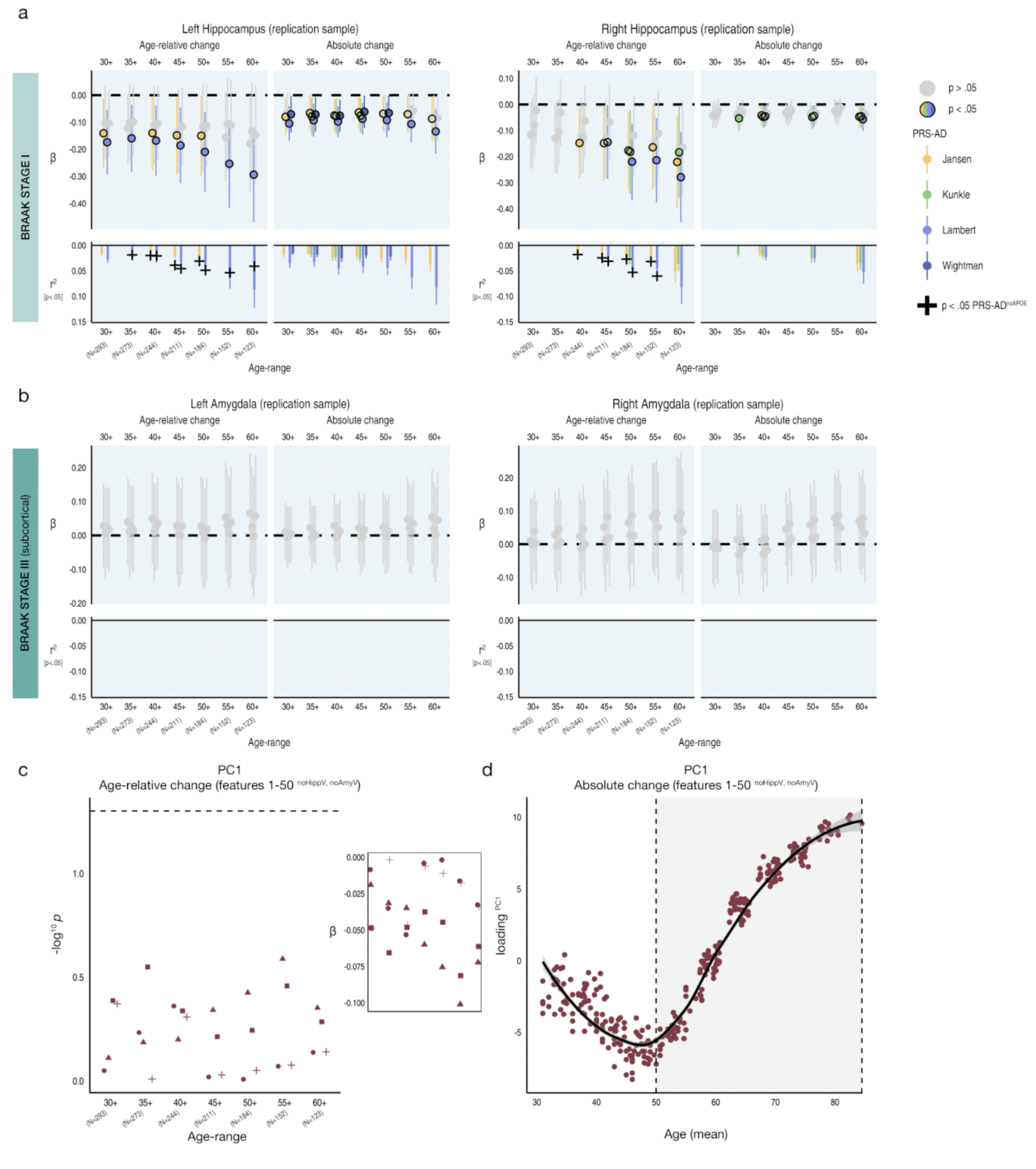
Replication. PRS-AD associations with age-relative and absolute change in an independent adult lifespan sample (Lifebrain replication sample), using the four GWAS-derived scores, for progressively older age-ranges to ensure capture of ageing-specific effects (i.e., moving from left to right on the X-axis, the leftmost age-range represents the association across the full adult lifespan on average [30-88 years; N=293], whereas the rightmost age-range shows the associations tested in only the oldest adults [60-88 years]). Univariate associations were tested for **a** left and right hippocampus, and **b** left and right amygdala. Significant associations at *p* < .05 are depicted in colour (upper panels). Here, partial r^2^ of PRS-AD is shown for all associations that were significant at *p* < .05 (lower panels). Where the association was significant (*p* < .05 [uncorrected]), we retested the association after removing *APOE* (PRS-AD^noAPOE^). Partial r^2^ of PRS-AD^noAPOE^ is depicted by a black cross if the association remained significant (*p* < .05). **c** Multivariate PRS-AD association tests using the principal component of age-relative change across the top 50 brain features with accelerated change in AD (excluding hippocampal and amygdala volumes; PC1^relChange^; as in Fig 4A-B). Datapoints show (-log10) p-values for the association with PC1^relChange^, tested at progressively older age-ranges, for all four scores. Smaller plot shows the standardized Beta values as a function of age-range (Betas inversed to be negative due to the non-directional nature of PCA). Dashed line indicates p = .05. **d** The principal component of absolute change across the top 50 brain features with accelerated change in AD (excluding hippocampal and amygdala volumes; maroon bar in Fig 4A), plotted as a function of mean age across timepoints. Accelerated brain change in AD-accelerated features was evident around age 50-60. Note that since the y-axis represents change, the slope of the curve represents acceleration (see also SI Fig. 10). Error bars depict 95% CI.

### Memory change analysis

Finally, in the LCBC healthy adult lifespan discovery sample, we used the association between the principal component of age-relative change across the first 50 AD-accelerated features – here including hippocampal and amygdala volumes (PC1^relChange1-50^) – and the principal component across the four PRS-AD scores (PC1^PRS-AD^; explaining 87%) to separate individuals into discrete groups, representing the conjunction of brain change and genetic risk factors. We hypothesized that high PRS-AD individuals also showing more age-relative change in AD-accelerated features would exhibit more longitudinal memory decline (pink quadrant 4 in Fig. 6D; Methods). Akin to the brain analysis, memory-change estimates were derived via the individual-specific random slopes in a GAMM of age, and we used longitudinal memory observations from the full adult lifespan sample to optimize memory-change estimates in the subset of participants that also had genetic data (Methods). Fig. 6A-C shows the longitudinal lifespan trajectory, and individual-specific degree of absolute and age-relative change in memory performance on the California Verbal Learning Test (CVLT; PC1 across subtests). Absolute memory change was predominantly negative, with memory decline occurring gradually across the adult lifespan and accelerating around the mid ∼60s (Fig. 6B; though we also observed a trend towards steeper slopes in mid-life prior to this; SI Fig. 11). As hypothesized, genetically exposed individuals also high on a multivariate marker of age-relative brain change (PC1^relChange1-50^) showed significantly more age-relative (*p* = .01) and absolute memory decline (*p* = .003) on average across the adult lifespan, compared to high PRS-AD individuals with less relative brain change. These group differences in memory change were not driven by differences in *APOE-ε4* carriership (Fig. 6E-F; main models corrected for carriership, mean age, sex, N timepoints, interval between first and last timepoint), and persisted in alternative models controlling for the number of *APOE-ε4* alleles (*p* = .009; *p* = .003) and baseline memory performance (*p* = .008; *p* = .002). In the main model, we also observed a significant difference in absolute memory change between the high PRS-AD-high brain change group and the low PRS-AD-low brain change group (*p* = .026; Fig. 6E). Finally, the reported group differences in memory-change persisted when correcting for differences in genetic risk (PC1^PRS-AD^) but not for differences in multivariate brain change (SI Fig. 12). These data suggest the conjunction of risk markers – a multivariate marker of change in AD-vulnerable features and known PRS-AD – helped identify a subset of comparatively high-risk individuals showing more longitudinal memory decline in healthy adult lifespan data (30-89 years).

**FIGURE 6.**
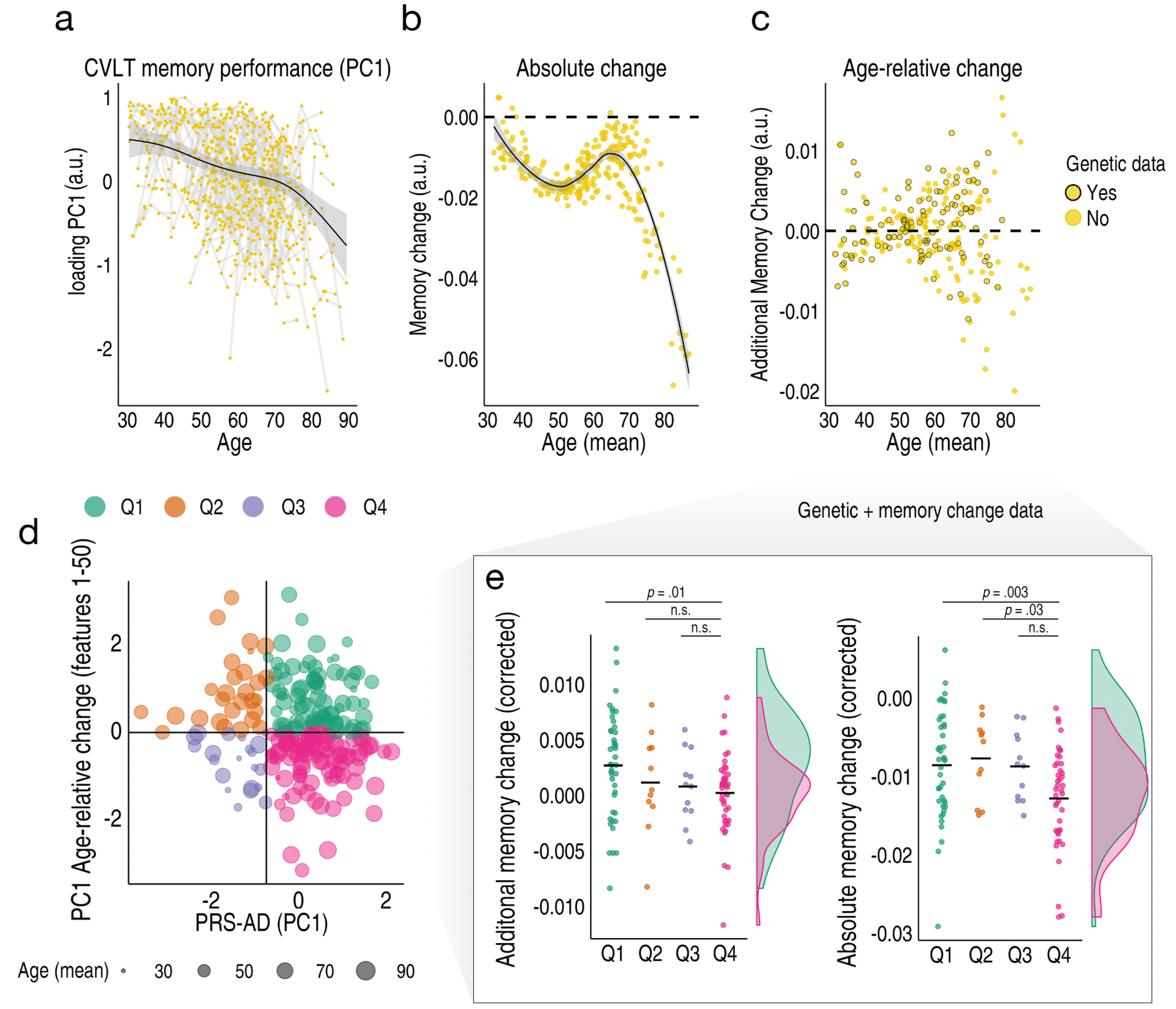
Longitudinal memory change analyses. Exclusively longitudinal data was used to estimate individual-specific age-relative and absolute change in CVLT task performance (PC1 across subtests), modelling the adult lifespan trajectories using GAMMs with random individual-specific slopes. **a** Adult lifespan trajectory analysis for CVLT memory performance from 30-89 years. Lines connect longitudinal observations per participant. **b** Absolute memory change per individual (datapoints) in CVLT task performance plotted as a function of their mean age across timepoints. **c** Estimated age-relative change per individual in CVLT task performance (individual-specific slopes). For each participant with memory change data, black stroke indicates whether or not genetic data was available. **d** The association between the principal component across the four PRS-AD scores and the principal component of age-relative change across the first 50 ADNI-derived features (listed in Fig. 4A) was used to define four quadrant-groups representing the conjunction of brain and genetic risk factors. **e** Memory change for individuals with both memory change and genetic data within the quadrant groups. Individuals at higher PRS-AD who also exhibited more age-relative brain change (pink) in AD-accelerated features showed significantly more age-relative (left plot) and absolute (right plot) change in memory performance across the healthy adult lifespan, relative to high PRS-AD individuals estimated to show less relative brain change (distributions visualized for these two groups; datapoints corrected for covariates including mean age and *APOE-e4* carriership [Methods]).

## Discussion

Genetic AD risk is robustly associated with the slope of brain ageing in healthy adults. Specifically, we found healthy individuals changing faster than expected for their age in early Braak Stage regions – bilateral hippocampus, amygdala, and right entorhinal cortex – are at significantly enhanced genetic AD risk, and these polygenic associations extend beyond the risk conferred by *APOE* alone. We also found that multivariate change across many AD-accelerated brain features can be used to detect PRS-AD associations with faster-than-expected brain ageing in healthy adults, and demonstrate that accelerated change in AD features is evident in most healthy individuals over age ∼50. Furthermore, we find that ML models trained on longitudinal AD patient-control data can be directly applied to healthy adult lifespan data and the prediction relates to PRS-AD in healthy adults. Finally, high PRS-AD individuals showing faster-than-expected brain change exhibited more longitudinal memory decline compared to high PRS-AD individuals with less brain change, on average across the healthy adult lifespan (30-89 years), and independent of *APOE-ε4*. Thus, the conjunction of our novel multivariate brain change marker and known PRS-AD found a subset of individuals exhibiting more memory decline across the healthy adult lifespan.

### Age-relative brain change across the adult lifespan associates with genetic AD risk Univariate analyses: apriori ROI’s

Univariate analyses using change in early Braak stage regions consistently revealed significant PRS-AD associations in healthy adults, illustrating accelerated brain ageing in genetically at-risk individuals. The clearest genetic effects upon faster brain ageing were in bilateral hippocampi; healthy individuals at higher genetic AD risk lose hippocampal volume faster than their age would predict – observed consistently using all four scores. Particularly for left hippocampus, the association often remained after discounting *APOE*, suggesting differences in left hippocampal loss also arise from genetic factors beyond *APOE*. However, we also observed PRS-AD^noAPOE^ associations with right hippocampal change, and also confirmed these in independent data. Shrinkage of the hippocampus – a critical structure underpinning episodic memory and spatial navigation operations – is a well-known AD risk marker in patient populations^9,32,54^, with atrophy rates predicting clinical conversion^55^. However, most studies in healthy adults have not linked genetic AD risk to hippocampal change^39,56^ or find the slope of hippocampal age trajectories does not differ as a function of genetic AD risk^2,20–26^ – including in large adult lifespan samples^24,25^ and our previous report in overlapping data^2^. And since AD risk genes influence hippocampal differences early in life^2,29,30^, cross-sectional findings in healthy older adults^27,28,57^ cannot attribute genetic effects to accelerated brain ageing^58^. By specifically isolating within-individual genetic effects on accelerated brain ageing, the present study confirms AD risk genes also influence normal variation in hippocampal change rates in healthy adults.

This agrees with a study by Harrison et al.^36^ finding a longitudinal relationship between hippocampal change and PRS-AD in older adults. Notably, however, that study recruited individuals with memory complaints and a family AD history via memory clinics. In contrast, our sample comprised healthy adults in longitudinal studies which are well-established to be biased toward maintaining high performers^59^. It also agrees with a study finding more hippocampal atrophy in healthy older *APOE-ε4* carriers^38^. However, we also found AD risk SNP’s beyond *APOE* predict hippocampal ageing trajectories in healthy adults, which to our knowledge has not been shown. Previously, we did not find consistent evidence PRS-AD or *APOE-ε4* alters the slope of hippocampal ageing, but found a group-level offset effect suggesting the difference between high- and low-risk individuals in hippocampal volume was as large at age ∼25 as at age ∼80^2^. However, that study primarily used a PRS-AD constructed with many more SNP’s (*p*<.05^60^), and did find some, albeit inconsistent, evidence for a slope effect using the same SNP association p-value as here. Here, by taking an individual-centric approach to estimate change trajectories, we found genome-wide significant SNPs could explain up to ∼13% variance in hippocampal change rates (effect sizes after discounting *APOE* were smaller; ∼5%; Fig. 1E-F). This purely longitudinal marker of *relative* brain ageing consistently excelled, exhibiting stronger relationships to PRS-AD than absolute change that were detectable over wider age-spans. The data also indicated PRS-AD-change associations were not driven only by the oldest adults, though older adults likely contributed more of the individual differences in brain change signal (SI. Fig. 12), in line with the observed tendency towards stronger genetic effects upon slopes in older individuals, and theories positing genetic effects become amplified in old age when neural resources are depleted^61^.

PRS-AD also linked with accelerated loss in right entorhinal cortex (stage I) and bilateral amygdala (stage III). This also agrees with Harrison et al.^36^, wherein entorhinal change was related to a PRS-AD (*APOE* inclusive) in older adults with memory complaints, and may also fit with a recent cross-sectional study finding right entorhinal cortex exhibits amongst the largest structural differences in older *APOE-ε4* carriers^28^. However, we also found evidence PRS-AD-entorhinal change associations extend beyond *APOE*. Similarly, accelerated amygdala loss was associated with PRS-AD in healthy adults, and we found evidence SNP’s beyond *APOE* influence left amygdala change trajectories. These data contradict a recent GWAS finding the effect of *APOE* upon amygdala and hippocampal slopes, with increasing influence of the *APOE*-indexing SNP (*rs429358*) with age, disappeared after accounting for disease in a heavily patient-derived sample^39^, suggesting *APOE*-mediated slope differences were driven by patients. To our knowledge, we are the first to document accelerated amygdala decline in healthy adults harbouring more AD risk variants. Still, while amygdala effects were clear in the discovery sample – currently the most densely sampled MRI dataset for longitudinal lifespan follow-up – these did not replicate in an independent sample with less follow-up, hence this awaits replication. Regardless, in healthy ageing as in AD, medial temporal lobe structures exhibit early vulnerability to structural loss^5^, highest expression of top AD risk genes (e.g., *APOE, BIN1, CLU*^62–64^), and we find PRS-AD influences accelerated change in these structures in healthy adults. Speculatively, faster atrophy rates may co-occur with faster tau accumulation, possibly consistent with higher tau in risk-allele carriers^64,65^. Critical questions concern what mechanisms underlie the shared vulnerability of these structures to lifespan influences and AD, which in the presence of AD risk genes speed up normal age-related neurodegeneration. One candidate shared characteristic may be a high degree of plasticity^66–68^.

### Multivariate analyses: data-driven features exhibiting accelerated change in AD

Through empirically delineating brain features with accelerated change in AD, we found that accelerated brain ageing across many combinations of AD-accelerated brain features relates to PRS-AD in healthy adults. Furthermore, we observed replicable evidence that almost everyone above age ∼50 is on an accelerated trajectory of neurodegenerative ageing in features wherein change reliably separates AD patients from controls, consistent with work documenting overlapping mean atrophy patterns in ageing and AD^4,5,14^. These individualized data suggest that neurodegeneration occurs along a continuum from healthy ageing to AD. Furthermore, since it is unlikely that most healthy adults in both samples here would be amyloid positive, this may run counter to the amyloid cascade hypothesis, which posits plaque build-up as an initial triggering event for subsequent neurodegeneration^69–71^. Likely, our unique approach to link AD changes to normal ageing benefitted from using multivariate analyses across change data in healthy adults. We also found that ML models trained on longitudinal change in AD can be applied to healthy adult lifespan data and the prediction relates to PRS-AD. This seemed to work best when the model was trained on estimates of change conditional on age (SI Fig. 5), likely because this places often extreme change values in AD on a scale more comparable across ages, and because modelling relative change in AD versus controls enables identification of features exhibiting a quantitative difference in change despite the presence of a similar qualitative pattern. That our patient-control groups were based on two extremes (consistently healthy versus becoming AD) only further emphasizes the difference lies more in degree than kind, as does the fact that our ML model also captured 100% of independent AD cases (Fig. 3). PRS-AD associations beyond Braak stages appeared largely though not entirely driven by *APOE* (Fig. 4; SI Fig. 5). Thus, our study yields new knowledge on the widespread impact of AD risk genes upon accelerated brain ageing in healthy adults, while highlighting that the border between neurodegeneration in ageing and AD is far from clear.

Of note, though PRS-AD effects were not entirely driven by allelic variation in *APOE*, PRS-AD^noAPOE^ associations were most evident using the genome-wide significant SNP’s/weightings reported by Jansen et al.^72^ or Lambert et al.^60^, suggesting these SNP sets beyond *APOE* better capture differences in brain ageing in healthy adults (in both samples; Figs. 1-2; Fig. 5). We also found no evidence including more SNP’s increased sensitivity to detect genetic effects upon healthy adult brain ageing, with or without APOE (SI Fig. 2), in line with studies in patients^73,74^. *APOE* accounted for much of the predictive power of PRS-AD, as associations typically disappeared or were attenuated using PRS-AD^noAPOE^. This fits with work finding PRS-AD associations with cognitive, lifestyle, and metabolic factors in healthy adults are largely driven by *APOE*^75^, and with data indicating limited utility of SNP’s beyond *APOE* to predict AD-relevant traits^18^.

### Memory change analysis

Individuals at higher genetic risk that also showed more brain ageing in AD-accelerated features exhibited more longitudinal memory decline across adult life (30-89 years). Hence, knowing an individual’s genetic risk in and of itself was insufficient, as it was not necessarily reflected in brain and cognitive outcomes. However, considered together with a multivariate marker of brain change, we found a subset of high PRS-AD individuals whose brain status over time was reflected in a greater drop-off in memory that was protracted across adult life (Fig. 6D-E). Moreover, the analyses suggested group differences in memory decline were more driven by brain change differences than by genetic differences. Hence, our change marker provided crucial information for detecting comparatively at-risk individuals in healthy adult lifespan data, beyond that provided by genetic risk alone. These results support and extend previous studies finding PRS-AD^43,44,46^ or *APOEε4*^45^ relates to longitudinal memory decline across adult life, and possibly shed light on why reported associations are often weak^43–46^ or absent^49^. They also underscore the need for follow-up data over extended age-spans when the goal is early prediction or prevention of AD. Future research should examine the biological and exposure-related factors that lead some high PRS-AD individuals to decline more in brain and memory where others remain resilient, as well as combine multivariate change with other biomarkers (e.g., tau, inflammation, or amyloid) as we move towards a future of individualized risk assessment.

Our study has several strengths. First, our longitudinal marker of individual-specific brain change circumvents the drawbacks of other approaches attempting to capture interindividual differences in brain ageing – such as brain age models^76^ – which do not necessarily relate to longitudinal change^77^. Second, we used the full breadth of the adult lifespan data to estimate individual-specific brain and memory change, each in a single model, using all longitudinal scans. This likely optimized the change estimates for all, including the subset with genetic data, likely in part due to improved age trajectory modelling from which one can subsequently estimate the deviation of an individual’s change trajectory. This is exemplified in SI Fig. 9, wherein we found PRS-AD-change associations in the same individuals in the BETULA study improved when their individual-specific slopes were estimated together with NESDA study data, compared to when estimated in BETULA data alone. Further, largely to ensure we were capturing ageing-specific processes at some point (see SI Fig. 1), we allowed the data to be increasingly comprised of only older individuals and repeatedly tested PRS-AD associations with change. As inferences based on significance are affected by arbitrary analysis choices, we took inspiration from multiverse methods to systematically define a defensible set of analysis choices to perform analyses across^51,52^. In our case, the principle arbitrary covariate was the age-range to test the association across, and the influence of this arbitrary choice on statistical significance is made clear in Fig. 2, Fig. 4B and Fig. 5, despite accounting for age- and time-related covariates. Adopting this approach, we could ensure capture of ageing-specific processes, document the stability of PRS-AD-change associations in healthy adults, and ensure the results were independent of a single arbitrary decision^51,52^, thus increasing their robustness.

There are also limitations. First, our approach disregards heterogeneity in ageing or AD-related atrophy; we considered all individuals obtaining an AD diagnosis over time as a single group, contrasting their average change against all consistently healthy individuals. For our purpose of delineating features with faster average change in AD, this was reasonable, as there may be a predominant AD atrophy pattern^78^ and it is this that overlaps with the average ageing pattern^5,6,42^. However, as there are known AD subtypes^78–80^, an important question is whether individual variability in AD atrophy presentation traces to heterogeneity in brain change in healthy adults. Second, as with most large-scale brain studies, we relied on FreeSurfer-derived measures. While these are well-validated and reliable^81–83^, it is possible measures such as entorhinal cortex may be less reliable^83^. Indeed, that we observed no PRS-AD associations with left entorhinal change was surprising, and possibly manual entorhinal tracing may have led to different results. Third, longitudinal lifespan studies inevitably culminate in unrepresentative samples comprised of a higher proportion of cognitively high-performers^59^. Since even in healthy adults we find variation in brain ageing slopes that maps onto AD-related genetic variation and memory outcomes, it is possible the population effect-sizes may be larger. Fourth, we used only structural MRI measures sensitive to detecting small changes in brain structure that ultimately form a continuous, lifelong process of change. Including additional imaging or biomarkers will help refine detection of AD-risk in healthy adults. Finally, we do not know which individuals included here will be diagnosed with AD later in life. While our analyses suggest one could assign differential transition probabilities to healthy individuals, only time and follow-up data will tell.

## Conclusion

In conclusion, brain ageing trajectories in healthy adults are robustly altered by the presence of AD risk genes, in many brain features, and beyond *APOE*. We show brain features most susceptible to faster deterioration in AD are on a trajectory of accelerated change from age ∼50 in healthy individuals, and that models trained on AD patients can be applied to adult lifespan data and the prediction relates to genetic AD risk in healthy adults. Finally, genetically at-risk individuals also high on a marker of brain change showed more adult lifespan memory decline, compared to genetically at-risk individuals with less brain change – suggesting our brain change marker enhanced the value of already knowing an individual’s genetic risk. AD risk genes are likely not AD-specific, but induce variation in the speed of the shared pattern of ageing- and AD-related neurodegeneration along a continuum in healthy adults. Our results call for a dimensional approach to late-onset AD as not being clearly distinct from normal brain ageing.

## Methods

### Samples

#### Age-relative change estimation

##### Adult lifespan discovery sample

After applying exclusion criteria (see below), an exclusively longitudinal adult lifespan sample (minimum two timepoints) comprising 1430 scans from 420 healthy individuals aged 30 to 89 years (248 females; mean age [SD] = 63.7 [14.4]; 2-7 timepoints [median = 3]; follow-up range = .15 - 11.1 years) was drawn from the Center for Lifespan Changes in Brain and Cognition database (LCBC; Department of Psychology, University of Oslo; see SI Note). Observations were collected across 3 scanners. Prior to participation, all individuals were screened via health and neuropsychological assessments, and the following exclusion criteria were applied across LCBC studies: evidence of neurodegenerative, neurologic or psychiatric disorders, use of medication known to affect the central nervous system (CNS), history of disease/ injury affecting CNS function, and MRI contraindications as assessed by a clinician. Additionally, to guard against including participants with incipient AD in our sample, we here excluded adults whose scores on the Mini Mental State Exam (MMSE)^84^ suggested longitudinal cognitive deficit with no later recovery (MMSE < 25 at their final timepoint; 2 participants; 4 scans), and adults aged 40+ whose scores on the Beck Depression Inventory (BDI)^85^ or Geriatric Depression Scale (GDS)^86^ suggested depression symptoms over time with no later recovery (BDI > 21 or GDS > 10 at their final timepoint; 7 participants; 32 scans). All LCBC studies were approved by the Norwegian Regional Committee for Medical and Health Research Ethics, complied with ethical regulations, and all participants provided informed consent.

##### Adult lifespan replication sample

To test replication, we used the two remaining longitudinal adult cohorts from the Lifebrain consortium that had up to three MRI timepoints available: the BETULA project^87^ and the Netherlands Study of Depression and Anxiety (NESDA) ^88^. BETULA participants underwent dementia assessment by a clinician using cognitive data and medical records, and those reporting neurological disorders (stroke, AD, other dementias, MS), or presenting with severe memory deficits or MRI contraindications were excluded. NESDA participants reporting neurological disorders (stroke, AD, other dementias, MS), or presenting with severe memory deficits or MRI contraindications were excluded. One extreme outlier in the change data of each sample was also detected and excluded here (see SI Fig. 8). In all, we collated the data from 449 scans from 182 individuals aged 31 - 88 from BETULA (mean age = 64.3 [11.9], 2-3 timepoints, follow-up = 3.5 – 7.7 years; 85 females), with 331 scans from 138 individuals from NESDA aged 30 - 65 (mean age = 45.1 [7.9], 2-3 timepoints, follow-up = 1 – 10 years; 91 females), into a single adult lifespan replication sample (SI Table 4). Although neurologically normal, 97 of the NESDA participants were diagnosed with a current or remitted depressive and/or anxiety disorder, whereas 41 had no history of mental health disorders.

#### Polygenic risk associations

To test associations with PRS-AD we used the subset of participants with both quality-controlled genetic data (European ancestry) and longitudinal change estimates, as estimated from the full adult lifespan models with all participants (also those without genetic data). For the discovery sample, 229 participants had genetic and brain change data. For the replication sample, 175 participants from BETULA and 118 from NESDA (92 diagnosed) had genetic and brain change data.

##### AD samples

We used exclusively longitudinal data from the Alzheimer’s Disease Neuroimaging Initiative (ADNI^89^), and the single-timepoint ADNI diagnosis (normal controls [NC]; mild cognitive impairment [MCI]; AD) to define two longitudinal groups based on final-timepoint diagnosis (2–9 timepoints): *NC-long* consisted of subjects classed as NC at every diagnosed timepoint; *AD-long* consisted of all subjects where the final diagnosed timepoint was AD^7^. After grouping, for subjects where scanner field strength changed over time (from 1.5T to 3T), we used observations from the scanner with the most timepoints (or where equal used the 3T scans). In all, *NC-long* consisted of 1680 scans from 372 subjects, and *AD-long* consisted of 2730 scans from 606 subjects (SI Table 2). The ADNI (PI: Michael W. Weiner, MD) was launched in 2003, with a goal of testing whether serial MRI can be used to measure the progression of MCI and early AD (see https://adni.loni.usc.edu/about/). An independent AD-control sample consisting of 107 scans from 39 *AD-long* subjects and 435 scans from 128 *NC-long* subjects was used for validation of ML models (AIBL dataset; data collected by the AIBL study group^90^; SI Fig. 4).

#### Genotyping and polygenic scores

In the LCBC dataset, buccal swab and saliva samples were collected for DNA extraction, followed by genome-wide genotyping using the Global Screening Array (Illumina, Inc., San Diego, CA). For a full description of genotyping, post-genotyping, and quality control and imputation methods applied to the genetic samples here, see^2,91,92^. We used the summary statistics from four previous large-scale GWAS of AD^60,93^ two of which included AD-by-proxy subjects based on parental status^72,94^. We then computed polygenic risk scores based on the genome-wide significant SNPs reported in each (*p* < 5 ×10^−8^), weighted by their allelic effect sizes. Prior to this, shared SNPs between each GWAS and our data were pruned to be nearly independent using PLINK^95^ with the following parameters --clump-p1 0.9999 --clump-p2 0.9999 –clump-r2 0.1 –clump-kb 500. The linkage disequilibrium structure was based on the European subpopulation of the 1000 Genomes Project Phase 3^96^. Because of the complexity of the major histocompatibility complex region (build hg19; chr6: 25,652,429-33,368,333), we removed SNPs in this region except the most significant one prior to pruning. We computed the four PRS-AD both with and without SNPs from the *APOE* region (build hg19; chr19: 44,909,011-45,912,650). We chose a genome-wide significant SNP threshold based on recent studies showing highest discrimination ability between patients and controls^73,74^. We also reasoned PRS’ constructed with more relaxed p-value thresholds will be less comparable across the four scores. As an exploratory analysis, we tested two other thresholds proposed to be optimal in patient-control data (p<10^−572^; p<0.1^97^). From the summary files, we removed SNPs not in the reference data, with minor allele frequencies <.05, or with low imputation scores. Genetic ancestry factors (GAFs) were computed using established principal components methods. For the discovery sample analyses, we used the first 10 as covariates in genetic analyses^98^. For genetic analyses in the combined Lifebrain replication sample, the first 4 were used as covariates (NESDA data was prepared using ENIGMA protocols requiring 4 GAFs^91^).

#### MRI acquisition and pre-processing

T1-weighted (T1w) anatomical scans from each MRI dataset (acquisition parameters in SI Table 5) were processed using FreeSurfer’s longitudinal stream^99^ (v.7.1 for LCBC, BETULA, ADNI and AIBL, v6.0 for NESDA), yielding a reconstructed cortex and subcortex for each participant and timepoint^100,101^. Data for the main discovery sample comprised T1w magnetization prepared rapid gradient echo (MPRAGE) sequences collected on 3 scanners at Oslo University Hospital; a 1.5T Avanto (599 scans), a 3T Skyra (769 scans), and a 3T Prisma (62 scans; Siemens Medical Solutions, Germany).

#### A priori ROIs

We first analyzed subcortical and cortical volumes for *a priori* defined ROI’s based on known AD vulnerability. These were based on the Braak staging scheme, initially defined using post-mortem measures of tau^50^ and subsequently applied to *in vivo* imaging^102^. Similar to others^102,103^, we used FreeSurfer regions from the aseg and Desikan-Killiany (DK) atlas^104^ that anatomically approximate the various stages (see https://ja-gustlab.neuro.berkeley.edu/s/Braak_ROI-3l2g.pdf). ROIs were constructed separately per hemisphere^7^. After initial analyses with our main hippocampal ROI’s – corresponding to Braak Stage II^50^ – we analyzed ROI’s corresponding to Stages I (entorhinal) and Stage III^50^, the latter we subdivided into a subcortical (amygdala) and a composite cortical ROI (parahippocampal, fusiform, lingual).

#### Data-driven ROIs

To empirically derive brain features with accelerated change in AD, we used machine learning in ADNI data (below) on a total of 364 features from the aseg and DK atlas^104^, comprising measures of cortical volume, area, thickness, grey matter/white matter contrast, subcortical volume and intensity (Fig. 3). This set of 364 features was also extracted and modelled within the discovery and replication samples.

## Statistical analysis

### Age-relative brain change across the adult lifespan

We used Generalized Additive Mixed Models (GAMMs, gamm4 v 0.2-6^105^) to estimate age models for each of the 364 brain features, fitting a nonlinear term for age (corrected for sex, scanner, and intracranial volume, knots = 8). We specified random intercepts and slopes for each participant. This enabled fitting an individual-specific linear model (level and slope) across all of their timepoints, to estimate how each person’s slope as a function of age deviates from the average nonlinear estimation. For an age model of e.g., hippocampus, random slopes are interpretable as the extent of additional (or reduced) hippocampal change an individual exhibits relative to the predicted change given their age (taking other covariates into consideration). Hence, we refer to this as an estimate of “age-relative change”. To partition unique variance associated with individual-specific slope, the estimation requires that a number of participants have three or more timepoints, although estimates are also produced for participants with fewer, but then are drawn from a population distribution more skewed towards the sample mean ^106^. This estimation is equivalent to estimating factor scores, and as such is psychometrically superior to manual calculations of change. Absolute change was calculated by adding the random slopes to the first derivative of the GAMM average age trajectory.

### Polygenic risk associations

#### Univariate analysis: a priori ROI’s

For each of our *a priori* ROIs, we used the random slopes as response variable in linear models with a PRS-AD predictor and the following covariates: mean age (across timepoints), sex, N timepoints, interval between first and last timepoint, and 10 genetic PCs (GAFs). We tested the associations between PRS-AD (4 scores; tested separately) and age-relative change with progressively older age-ranges (i.e., 30-89, 35-89, 40-89 … 70-89). The reasons for this were threefold. First, because some brain features were estimated to have more negative individual-specific slopes in younger adults compared with middle-age (SI Fig. 1), we could not test the association across the entire age-range (30-89) and ensure we were capturing only ageing-specific processes. Second, it enabled assessing the stability of PRS-AD associations detectable in adult lifespan data (note that older age-ranges correspond to smaller sample sizes). Third, because empirical outcomes are influenced by arbitrary analysis decisions, we took inspiration from multiverse methods that attempt to reduce such bias by testing associations across a set of theoretically justified alternatives ^51,52^. We also tested each association with absolute change, and False Discovery Rate (FDR) correction was applied across all 576 PRS-AD tests (8 structures × 4 scores × 9 age-ranges × 2 change metrics; significance considered at *p*[FDR] < .05). For surviving PRS-AD associations, we tested whether the FDR-corrected association including *APOE* remained significant at *p* < .05 using PRS-AD^noAPOE^, and determined whether the number of significant hits exceeded the 5% false positive rate per structure. We also ran post-hoc tests to confirm that the PRS-AD-change estimates became more negative as the age subset steadily comprised only older individuals (see Fig. 1E-F). Here, we used the pre-computed beta estimates from all PRS-AD-change models (age-relative and absolute; all four scores) as response variable, and the age-range as predictor (coded 0-8), and tested the linear effect of age-range upon the PRS-AD beta estimates (main effect across change models). The observed coefficient thus represents the strengthening of the negative PRS-AD-change association for each increasing age subset. Next, we permuted the empirical p-value for this observed association, by generating a null distribution across 1000 random permutations of the age variable (mean age) in the PRS-AD change associations, then recalculating the effect of age-range (randomized) upon the PRS-AD beta estimates.

### Multivariate analyses: data-driven features exhibiting accelerated change in AD

#### Machine learning model in AD

We repeated the procedure to estimate age-relative change in ADNI data, fitting a GAMM of age across *NC-long* and *AD-long* groups (Fig. 3A; covariates: sex, field strength). To guard against overfitting the age trajectories and account for the roughly three-decade drop in age coverage in the AD datasets (SI Table 2), we reduced the number of knots in the GAMM to 5. Next, we ran machine learning binary classification with XGBoost (https://xgboost.readthedocs.io^53^), using the random individual-specific slopes (age-relative change) across all 364 features as input. Hyperparameters were chosen using 10-fold cross validation across 500 random combinations of the following possible parameter values: nrounds (100 – 600, step = 50), eta (0.01, 0.05, 0.1, 0.15, 0.2), max_depth (2-8, step = 1), gamma (0.5 – 1.5, step = 0.5), min_child_weight (1 – 4, step = 1). To reduce the risk of overfitting to the training data and increase generalizability, we selected the final hyperparameters based on the mean AUC obtained across the 500 iterations of 10-fold cross-validation, where each iteration logged the maximum AUC achieved across folds (final hyperparameters: nrounds = 500, eta = 0.2, max_depth = 5, gamma = 1, min_child_weight = 1). This approach ensures a more robust and stable estimate of model performance across diverse data subsets while also avoiding potential overfitting to a single hyperparameter combination. For comparison, we also computed a classification model using absolute brain change as input following the same procedure (hyperparameters: nrounds = 600, eta = 0.01, max_depth = 7, gamma = 0.5, min_child_weight = 2). Model performance was evaluated in AIBL data (Fig. 3; SI Fig. 4).

#### Application to healthy adult lifespan data

First, we extracted the feature matrix to derive a list of brain features important for classifying *AD-long* from *NC-long* individuals based on age-relative change in ADNI. Then, in the LCBC healthy adult lifespan discovery sample, we calculated the principal component of age-relative change (PC1^relChange^) across the top 50 features, not including hippocampal and amygdala volumes (to ensure these did not drive the effect). We then used PC1^relChange^ to test for PRS-AD associations with change in our healthy adult lifespan sample, at progressively older age-ranges, for all four scores. Next, we aimed to ensure the observed multivariate associations were not disproportionately driven by one or a few brain features. To do this, we first calculated the age at which absolute brain change accelerates, reasoning analyses within this age-range would give maximal chance of detecting PRS-AD effects upon individual ageing trajectories. Here, we took the principal component of absolute change across the same set of features (PC1^absChange^), plotted as a function of mean age. Then, within the 50-89 years age-range (Fig. 4C), we ran a sliding window PCA, iteratively calculating PC1 across 20 features with a step size of 3, across the first ∼100 features (complete windows of 20 up to 98 features; 27 windows), and tested PC1 associations with PRS-AD within each window. FDR-correction was applied across all 144 PRS-AD tests in this analysis, and surviving associations were tested with PRS-AD^noAPOE^.

As a final proof-of-principle, we applied the weights from the binary classification procedure in AD-control data directly to the healthy adult lifespan data (i.e., LCBC as test data). This prediction uses information from the weights of all 364 features. Here, the dependent variable was calculated as log[p/(1-p)], where p is the model-implied probability of having AD (probAD^relChange^). The aim of this was not to classify healthy individuals as AD or not, but rather test our hypothesis that the learned model weights would nevertheless prove useful, and would relate to PRS-AD in healthy adult lifespan data. We also tested whether predictions derived from the ML model based on absolute change were related to PRS-AD. Again, FDR-correction was applied across all 72 PRS-AD tests in this analysis, and surviving associations were tested with PRS-AD^noAPOE^.

#### Replication analysis

We first ran a GAMM separately in each of the replication cohorts, revealing a strong outlier for each in the hippocampal change data (-7.4SD in BETULA; +5.5SD in NESDA; see SI Fig. 8). Then, we collated the data and ran a GAMM comparable to the main analysis (scanner covariate indexed study cohort), estimated the random slopes, and excluded these two outliers (SI Fig. 8). Similar to the main analysis, we expected including as many longitudinal observations as possible in the GAMM would optimize the change estimates for all. Testing this assumption post-hoc, we found that in the same individuals with genetic data from BETULA, beta estimates with left hippocampal change were significantly lower when their random slopes were estimated together with NESDA data, relative to only using BETULA data (*p* = .009; SI Fig. 9). To reduce the number of tests, we tested PRS-AD associations with change in hippocampus and amygdala, and with PC1^relChange^ (top 50 AD-accelerated features excluding hippocampal and amygdala volumes). PRS-AD models matched the discovery sample, except for an added cohort covariate. We tested the model at progressively older age-ranges for all four scores (here until a lower age-bound of 60, above which the sample was comprised entirely of BETULA subjects). Where the association was significant (p < .05 [uncorrected]), we tested whether it remained significant with PRS-AD^noAPOE^. We considered it a replication where the number of significant tests per structure exceeded the 5% false positive rate. Lastly, we assessed whether the trajectory of accelerated brain ageing in AD features mirrored the discovery sample (i.e., modelled PC1^absChange^ as a function of mean age).

#### Memory change analysis

Finally, we tested differences in memory change between groups of individuals defined by the conjunction of brain and genetic risk markers. We hypothesized higher PRS-AD individuals also high on a multivariate marker of brain change would show more memory decline across the adult lifespan. This analysis proceeded in two parts. First, we took the principal component across the four PRS-AD scores (PC1^PRS-AD^; explaining 87%), and used the association between PC1^PRS-AD^ and the principal component across the first 50 AD-accelerated features (here including hippocampal and amygdala volumes), to divide individuals into quadrant groups (Fig. 6D; pink group denotes individuals high on both risk factors). Second, from the full adult lifespan discovery sample described above (N = 420; scans = 1430), we identified those with observations on the California Verbal Learning Test (CVLT)^107^. Of these, we discarded individuals with non-usable memory data (due to being part of on-off memory training projects at LCBC; see SI Note 1 for information on the projects that comprised the LCBC sample). In the resulting data (713 observations from 267 individuals), we took the principal component across the three main CVLT subtests (learning, immediate, and delayed free recall; scaled) to index general memory, expressed the loadings as a proportion of the maximum loading, and kept only those with longitudinal memory observations (707 observations from 261 individuals). Then, we ran a GAMM of age on Memory (sex corrected, knots = 8). Akin to the brain analysis, age-relative memory change was estimated via the random slopes, and absolute memory change was calculated by adding the slopes to the first derivative of the GAMM average age trajectory. Having estimated memory change using as many longitudinal CVLT observations as possible – 108 individuals had both memory change and genetic data (i.e., were included in the quadrant-groups). Finally, we tested our hypothesis that the high brain change-high PRS-AD group would exhibit more adult lifespan memory decline, setting this group to the intercept, in linear models of quadrant-group on memory change, correcting for group differences in mean age, sex, N timepoints, interval between first and last timepoint, and *APOE-ε4* carriership (main model). These were tested using both age-relative and absolute memory change. Alternative models correcting for the number of *APOE-ε4* alleles, baseline memory, PC1^PRS-AD^, and PC1^relChange1-50^, were also run.

## Acknowledgements

*Scripts were run on the Colossus processing cluster at the University of Oslo, and on resources provided by UNINETT Sigma2 (project NN9769K). LCBC funding: grant 302854 (FRIPRO; to Y.W.), European Research Council under grants 283634, 725025 (to A.M.F.), and 313440 (to K.B.W.); Norwegian Research Council (to A.M.F. and K.B.W.) under grants 249931 (TOPPFORSK) and, The National Association for Public Health’s dementia research program, Norway (to A.M.F). The Lifebrain project is funded by the EU Horizon 2020 Grant: “Healthy minds 0–100 years: Optimising the use of European brain imaging cohorts (Lifebrain).” Grant agreement number: 732592. The Betula project was supported by a Scholar grant from Knut and Alice Wallenberg’s (KAW) foundation to L.N. The infrastructure for the NESDA study (www.nesda.nl) is funded through the Geestkracht program of the Netherlands Organisation for Health Research and Development (ZonMw, grant number 10-000‐1002) and financial contributions by participating universities and mental health care organizations (VU University Medical Center, GGZ inGeest, Leiden University Medical Center, Leiden University, GGZ Rivierduinen, University Medical Center Groningen, University of Groningen, Lentis, GGZ Friesland, GGZ Drenthe, Rob Giel Onderzoekscentrum). Some of the data used in the preparation of this article were obtained from the Alzheimer’s Disease Neuroimaging Initiative (ADNI) (National Institutes of Health Grant U01 AG024904) and DOD ADNI (Department of Defense award number W81XWH-12-2-0012). ADNI is funded by the National Institute on Aging, the National Institute of Biomedical Imaging and Bioengineering, and through generous contributions from the following: AbbVie, Alzheimer’s Association; Alzheimer’s Drug Discovery Foundation; Araclon Biotech; BioClinica, Inc*.; *Biogen; Bristol-Myers Squibb Company; CereSpir, Inc*.; *Cogstate; Eisai Inc*.; *Elan Pharmaceuticals, Inc*.; *Eli Lilly and Company; EuroImmun; F. Hoffmann-La Roche Ltd and its affiliated company Genentech, Inc*.; *Fujirebio; GE Healthcare; IXICO Ltd*.;*Janssen Alzheimer Immunotherapy Research & Development, LLC*.; *Johnson & Johnson Pharmaceutical Research & Development LLC*.; *Lumosity; Lundbeck; Merck & Co*., *Inc*.;*Meso Scale Diagnostics, LLC*.; *NeuroRx Research; Neurotrack Technologies; Novartis Pharmaceuticals Corporation; Pfizer Inc*.; *Piramal Imaging; Servier; Takeda Pharmaceutical Company; and Transition Therapeutics. The Canadian Institutes of Health Research is providing funds to support ADNI clinical sites in Canada. Private sector contributions are facilitated by the Foundation for the National Institutes of Health (www.fnih.org). The grantee organization is the Northern California Institute for Research and Education, and the study is coordinated by the Alzheimer’s Therapeutic Research Institute at the University of Southern California. ADNI data are disseminated by the Laboratory for Neuro Imaging at the University of Southern California. The ADNI researchers contributed data but did not participate in analysis or writing of this report. Some of the data used in the preparation of this article were obtained from the Australian Imaging Biomarkers and Lifestyle Flagship Study of Ageing (AIBL) funded by the Commonwealth Scientific and Industrial Research Organisation (CSIRO), which was made available at the ADNI database (www.loni.usc.edu/ADNI). The AIBL researchers contributed data but did not participate in analysis or writing of this report*.

