## Supplementary information for "Accelerated brain change in healthy adults is associated with genetic risk for Alzheimer’s disease and uncovers adult lifespan memory decline"

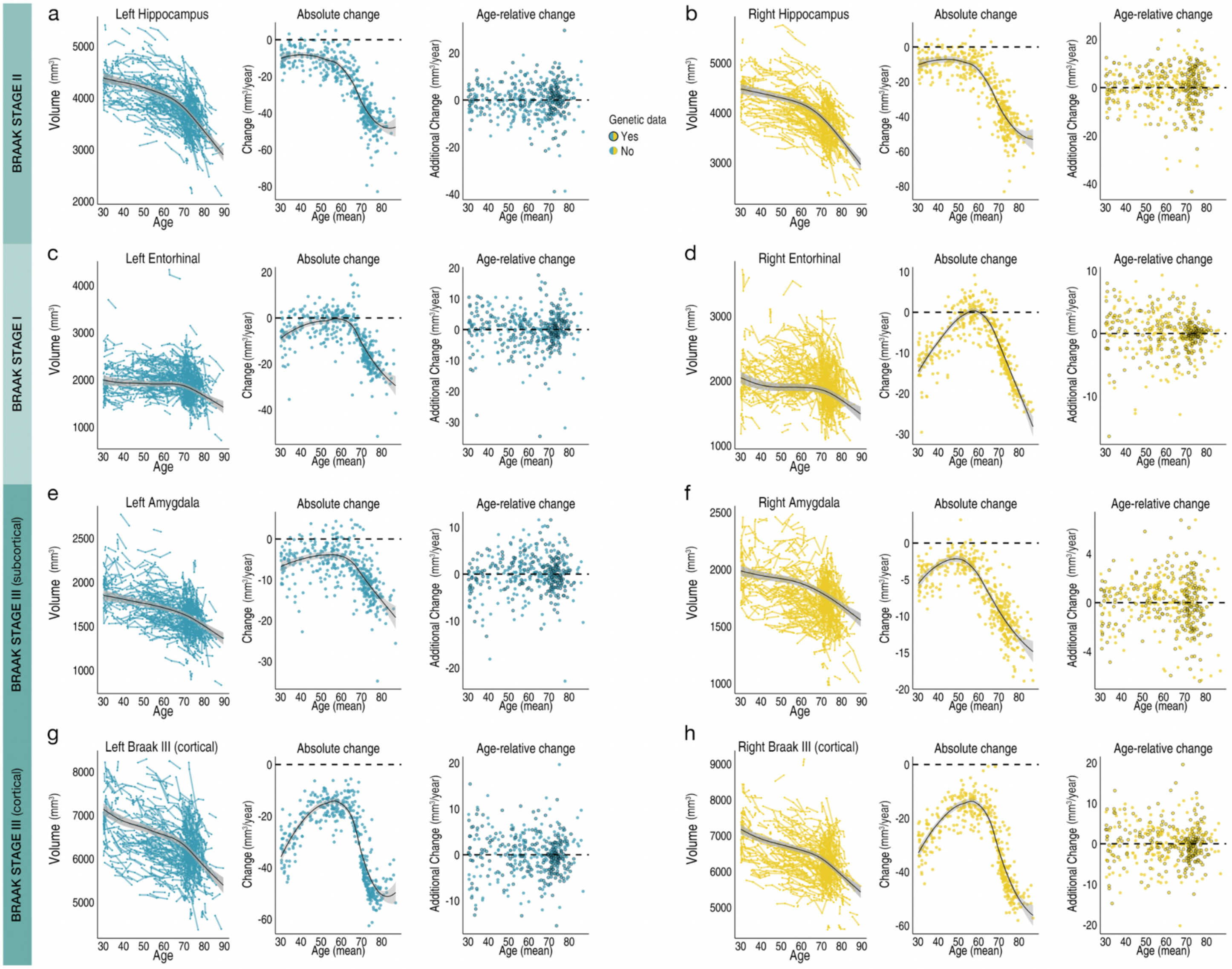

**SI FIGURE 1**

Exclusively longitudinal data was used to estimate individual-specific age-relative change in bilateral hippocampus (Braak stage II; **a** left, **b** right), entorhinal cortex (Braak stage I; **c,d**), amygdala (Braak stage III subcortical; **e,f**), and a Braak Stage III cortical region (**g,h**), modelling the adult lifespan trajectories using GAMMs with random individual-specific slopes. Leftmost plots in each: adult lifespan trajectory from 30-89 years (data corrected for sex and scanner, lines connect longitudinal observations). Middle plots: absolute change per individual (datapoints) as a function of their mean age across timepoints. Rightmost plots: estimated age-relative change per individual (i.e. individual-specific slopes) as a function of their mean age across timepoints. Black stroke indicates whether or not genetic data was available per participant and thus whether the datapoint was included in the PRS-AD association tests.

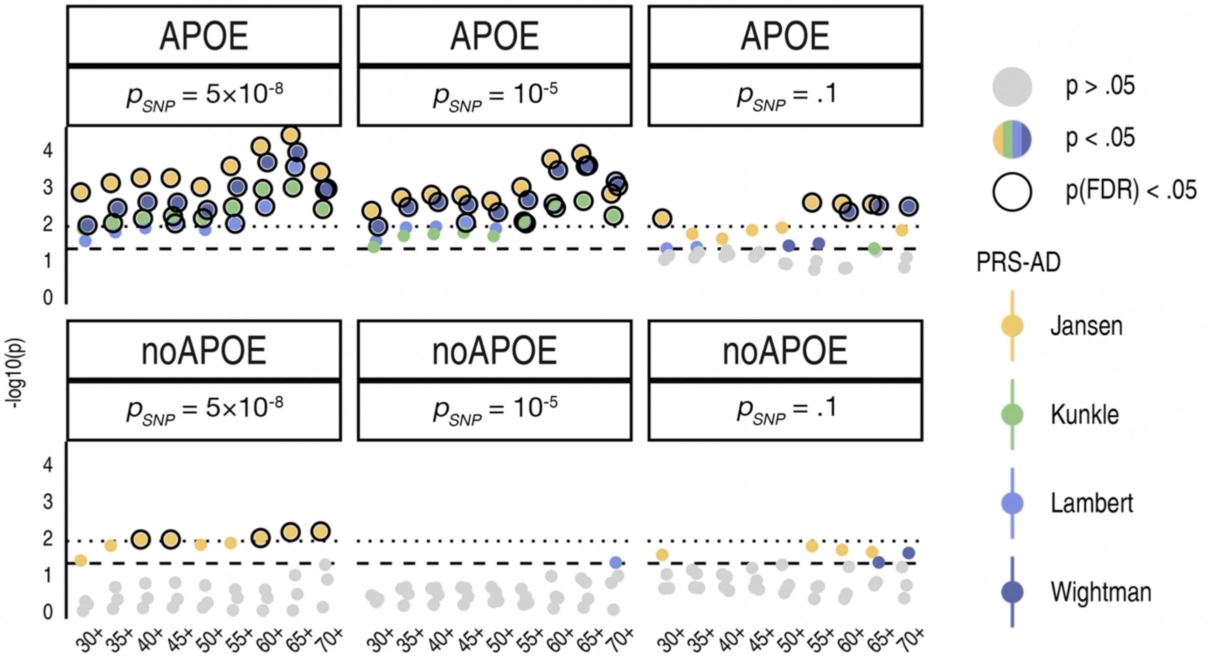

**SI FIGURE 2**

PRS-AD associations (-log10 p-values) with age-relative change in left hippocampus, with PRS-AD computed using two alternative p-value thresholds, and both with and without APOE. For comparison, the first column depicts the p-values using genome-wide significant SNPs as in main paper. Coloured points depict associations at p < .05 (uncorrected) and black outline depict associations surviving FDR-correction. As these tests are for illustrative purposes only and not independent of the main analysis, the FDR-correction level applied is the same as across all 576 PRS-AD tests reported in the univariate analysis in the main paper.

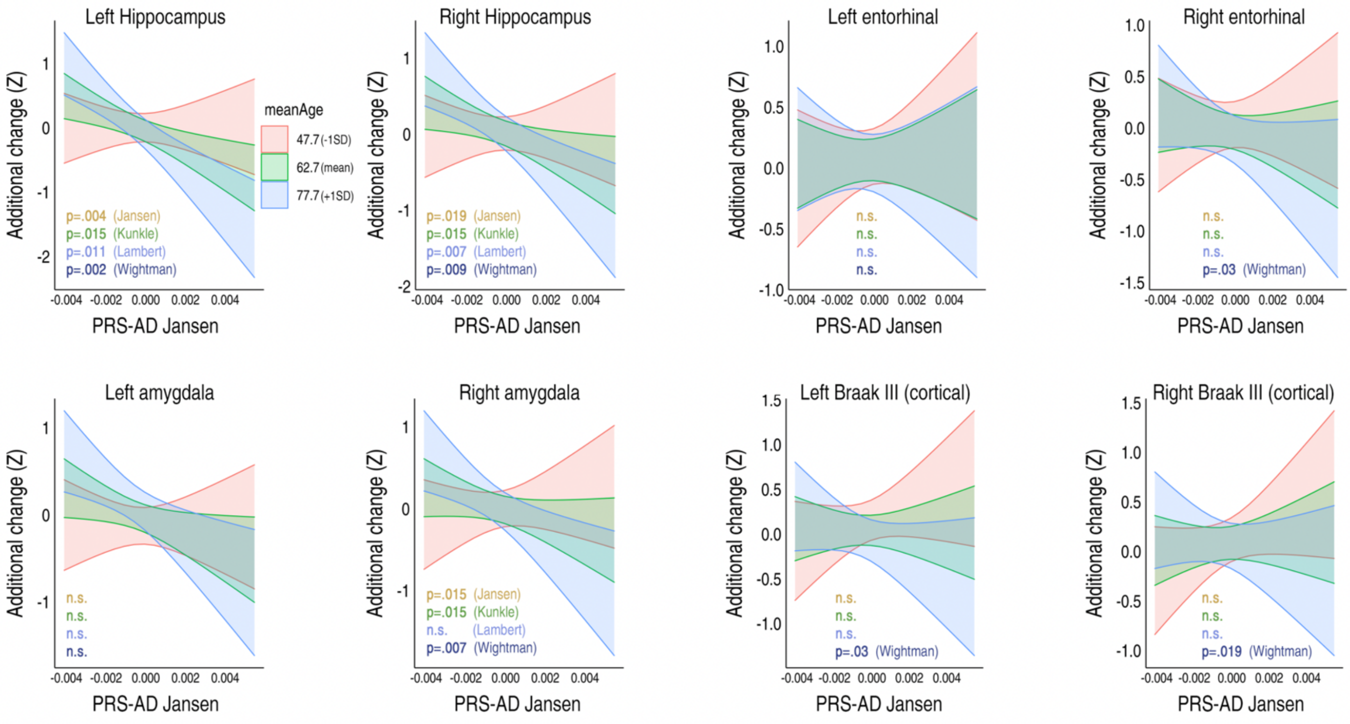

**SI FIGURE 3**

Results from alternative analyses dependent on power across the full age-range (30-89 years). Suggestively significant linear PRS-AD * mean age interactions upon age-relative change were found, indicative of steeper effects of PRS-AD upon age-relative change in older ages in these structures. However, none of the tests survived FDR-correction across the 32 tests performed in this analysis (see SI Table 1).

| stage | struct | score | effect | beta | stderr | t | p | df |
|---|---|---|---|---|---|---|---|---|
| Braak II | Left Hippocampus | Jansen | age_interaction | -0.21 | 0.07 | -2.92 | **0.00387** | 211 |
| Braak II | Left Hippocampus | Lambert | age_interaction | -0.18 | 0.07 | -2.58 | **0.01062** | 211 |
| Braak II | Left Hippocampus | Kunkle | age_interaction | -0.18 | 0.07 | -2.46 | **0.01487** | 211 |
| Braak II | Left Hippocampus | Wightman | age_interaction | -0.22 | 0.07 | -3.1 | **0.00219** | 211 |
| Braak II | Right Hippocampus | Jansen | age_interaction | -0.17 | 0.07 | -2.36 | **0.01925** | 211 |
| Braak II | Right Hippocampus | Lambert | age_interaction | -0.14 | 0.07 | -2.14 | **0.03351** | 211 |
| Braak II | Right Hippocampus | Kunkle | age_interaction | -0.19 | 0.07 | -2.74 | **0.00676** | 211 |
| Braak II | Right Hippocampus | Wightman | age_interaction | -0.18 | 0.07 | -2.63 | **0.00928** | 211 |
| Braak I | Left entorhinal | Jansen | age_interaction | -0.06 | 0.08 | -0.76 | 0.44868 | 211 |
| Braak I | Left entorhinal | Lambert | age_interaction | -0.05 | 0.07 | -0.66 | 0.50907 | 211 |
| Braak I | Left entorhinal | Kunkle | age_interaction | 0 | 0.07 | -0.01 | 0.98997 | 211 |
| Braak I | Left entorhinal | Wightman | age_interaction | -0.1 | 0.07 | -1.33 | 0.18490 | 211 |
| Braak I | Right entorhinal | Jansen | age_interaction | -0.1 | 0.07 | -1.37 | 0.17311 | 211 |
| Braak I | Right entorhinal | Lambert | age_interaction | -0.05 | 0.07 | -0.75 | 0.45180 | 211 |
| Braak I | Right entorhinal | Kunkle | age_interaction | -0.06 | 0.07 | -0.82 | 0.41544 | 211 |
| Braak I | Right entorhinal | Wightman | age_interaction | -0.16 | 0.07 | -2.21 | **0.02834** | 211 |
| Braak III (subcortical) | Left Amygdala | Jansen | age_interaction | -0.13 | 0.07 | -1.88 | 0.06098 | 211 |
| Braak III (subcortical) | Left Amygdala | Lambert | age_interaction | -0.1 | 0.06 | -1.54 | 0.12478 | 211 |
| Braak III (subcortical) | Left Amygdala | Kunkle | age_interaction | -0.11 | 0.07 | -1.57 | 0.11864 | 211 |
| Braak III (subcortical) | Left Amygdala | Wightman | age_interaction | -0.13 | 0.07 | -1.89 | 0.06026 | 211 |
| Braak III (subcortical) | Right Amygdala | Jansen | age_interaction | -0.18 | 0.07 | -2.46 | **0.01459** | 211 |
| Braak III (subcortical) | Right Amygdala | Lambert | age_interaction | -0.11 | 0.07 | -1.68 | 0.09410 | 211 |
| Braak III (subcortical) | Right Amygdala | Kunkle | age_interaction | -0.17 | 0.07 | -2.46 | **0.01485** | 211 |
| Braak III (subcortical) | Right Amygdala | Wightman | age_interaction | -0.19 | 0.07 | -2.7 | **0.00743** | 211 |
| Braak III (cortical) | Left temporal | Jansen | age_interaction | -0.14 | 0.07 | -1.89 | 0.06079 | 211 |
| Braak III (cortical) | Left temporal | Lambert | age_interaction | -0.07 | 0.07 | -1.07 | 0.28520 | 211 |
| Braak III (cortical) | Left temporal | Kunkle | age_interaction | -0.12 | 0.07 | -1.72 | 0.08707 | 211 |
| Braak III (cortical) | Left temporal | Wightman | age_interaction | -0.16 | 0.07 | -2.21 | **0.02817** | 211 |
| Braak III (cortical) | Right temporal | Jansen | age_interaction | -0.13 | 0.07 | -1.78 | 0.07726 | 211 |
| Braak III (cortical) | Right temporal | Lambert | age_interaction | -0.12 | 0.07 | -1.84 | 0.06663 | 211 |
| Braak III (cortical) | Right temporal | Kunkle | age_interaction | -0.13 | 0.07 | -1.83 | 0.06887 | 211 |
| Braak III (cortical) | Right temporal | Wightman | age_interaction | -0.16 | 0.07 | -2.37 | **0.01893** | 211 |

**SI TABLE 1**

Results from alternative analyses dependent on power across the full age-range (30-89 years; N = 229). Suggestively significant linear PRS-AD * mean age interactions (in bold) upon age-relative change were found, indicative of steeper effects of PRS-AD upon age-relative change in older ages in these structures. However, none of the tests survived FDR-correction across the 32 tests in this analysis.

| | | | | N Timepoints | | | | | | | | | | | | | Mean MMSE | | Mean Clinical Dementia Rating (CDR) | |
|---|---|---|---|---|---|---|---|---|---|---|---|---|---|---|---|---|---|---|---|---|
| Dataset | Group | N unique | N Obs | 2 | 3 | 4 | 5 | 6 | 7 | 8 | 9 | Mean Time Interval (SD) | Interval Range | Mean Age (SD) | Age-Range | Sex (f/m) | All Timepoints | Longitudinal Timepoints | All Timepoints | Longitudinal Timepoints |
| ADNI | NC-long | 372 | 1680 | 37 | 60 | 82 | 110 | 47 | 22 | 14 | 0 | 1.65 (1.47) | 0.05-6.66 | 75.4 (6.1) | 59.7 - 95 | 196 / 176 | 29.03 | 29.02 | 0.08 | 0.09 |
| | AD-long | 606 | 2730 | 55 | 86 | 197 | 147 | 53 | 35 | 21 | 12 | 1.5 (1.32) | 0.07-6.48 | 75.4 (7.3) | 55 - 92.9 | 257 / 349 | 23.94 | 23.53 | 3.95 | 4.24 |
| AIBL (test) | NC-long | 128 | 435 | 21 | 34 | 73 | | | | | | 2.9 (1.4) | 1.1-6.5 | 73 (7) | 60.5 - 90.2 | 65 / 63 | 28.95 | 28.95 | 0.02 | 0.01 |
| | AD-long | 39 | 107 | 17 | 15 | 7 | | | | | | 2.5 (1.1) | 1.3-5.2 | 74.9 (7.6) | 55 - 89.2 | 20 / 19 | 21.91 | 20.5 | 0.74 | 0.84 |

**SI TABLE 2**
Description of the longitudinal ADNI training data, and the longitudinal AIBL test data, by group status. Intervals are in years.

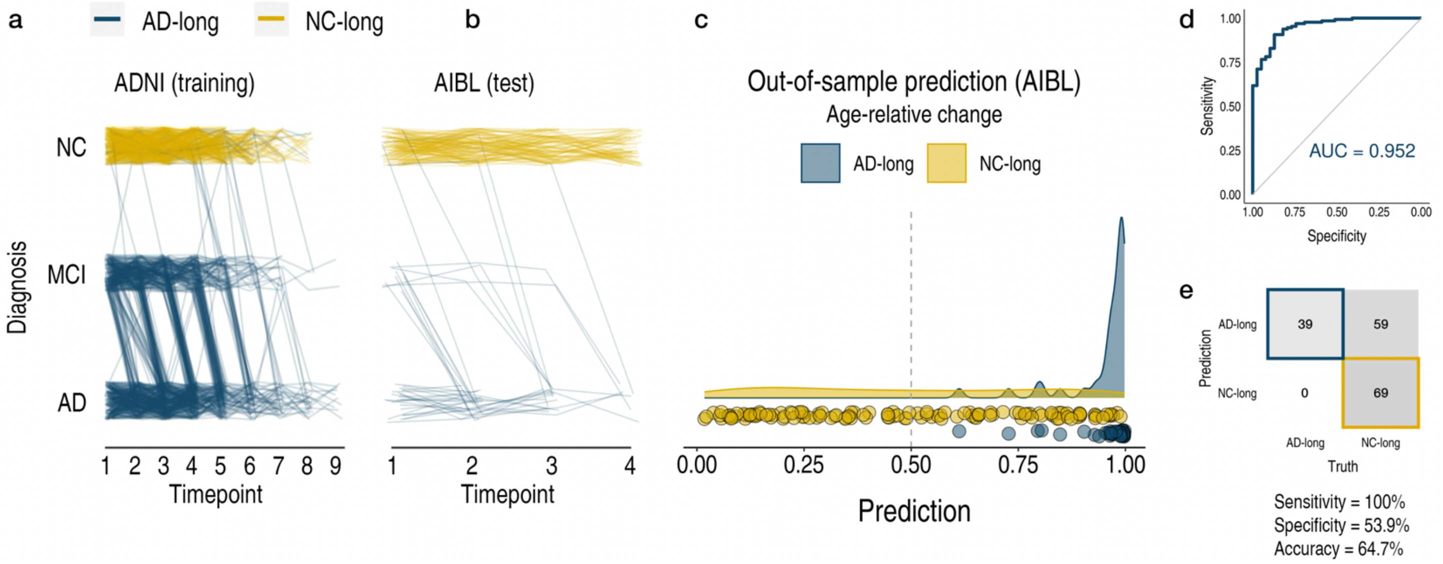

**SI FIGURE 4**
Visual description of the longitudinal ADNI training and AIBL test data. **a** (as in Fig. 3A in main paper) Longitudinal grouping in ADNI training data. X-axis denotes the scan observations across timepoints used in the final sample. Each line represents a participant and colour denotes longitudinal group membership. Single-timepoint ADNI diagnoses (Y-axis; NC normal controls, MCI mild cognitive impairment, AD Alzheimer's disease) were used to define two longitudinal groups of AD and NC individuals (AD-long; N = 606, obs = 2730; NC-long, N = 372; obs = 1680). *CN-long* individuals were classified as healthy at every timepoint whereas *AD-long* individuals were diagnosed with AD by their final timepoint (Methods). Note that trajectories of individuals that appear to end with a NC or MCI diagnosis nevertheless correspond to individuals with an AD diagnosis by their final timepoint, but either with no concurrent scan observation available, or no concurrent scan observation used due to scanner changes over time (Methods). Note also that individual trajectories seemingly reverting also correspond to individuals with an AD diagnosis by their final timepoint. **b** Longitudinal grouping in the independent AIBL test data. **c-e** Out-of-sample prediction for the binary classifier (AIBL) including receiver operator curve (d), confusion matrix and performance metrics (e).

| stage | struct | score | effect | beta | stderr | t | p | df |
|---|---|---|---|---|---|---|---|---|
| - | $PC1^{relChange}$ | Jansen | age_interaction | 0.23 | 0.07 | 3.21 | * **0.00154** | 211 |
| - | $PC1^{relChange}$ | AD | age_interaction | 0.19 | 0.07 | 2.74 | * **0.00664** | 211 |
| - | $PC1^{relChange}$ | Kunkle | age_interaction | 0.22 | 0.07 | 3.07 | * **0.00243** | 211 |
| - | $PC1^{relChange}$ | Wightman | age_interaction | 0.22 | 0.07 | 3.2 | * **0.00159** | 211 |

**SI TABLE 3**

Results from alternative analyses dependent upon power across the full age-range (30-89 years; N = 229). FDR-corrected significant linear PRS-AD * mean age interactions (in bolded denoted with *) upon upon $PC1^{relChange}$ were found for all four GWAS-derived scores, indicative of steeper effects of PRS-AD upon multivariate age-relative change across AD-accelerated features in older ages. Note that hippocampal and amygdala volumes were not included in $PC1^{relChange}$ to ensure these did not drive the multivariate effect (see Fig. 4A). FDR-correction was applied across all 4 tests performed in this analysis.

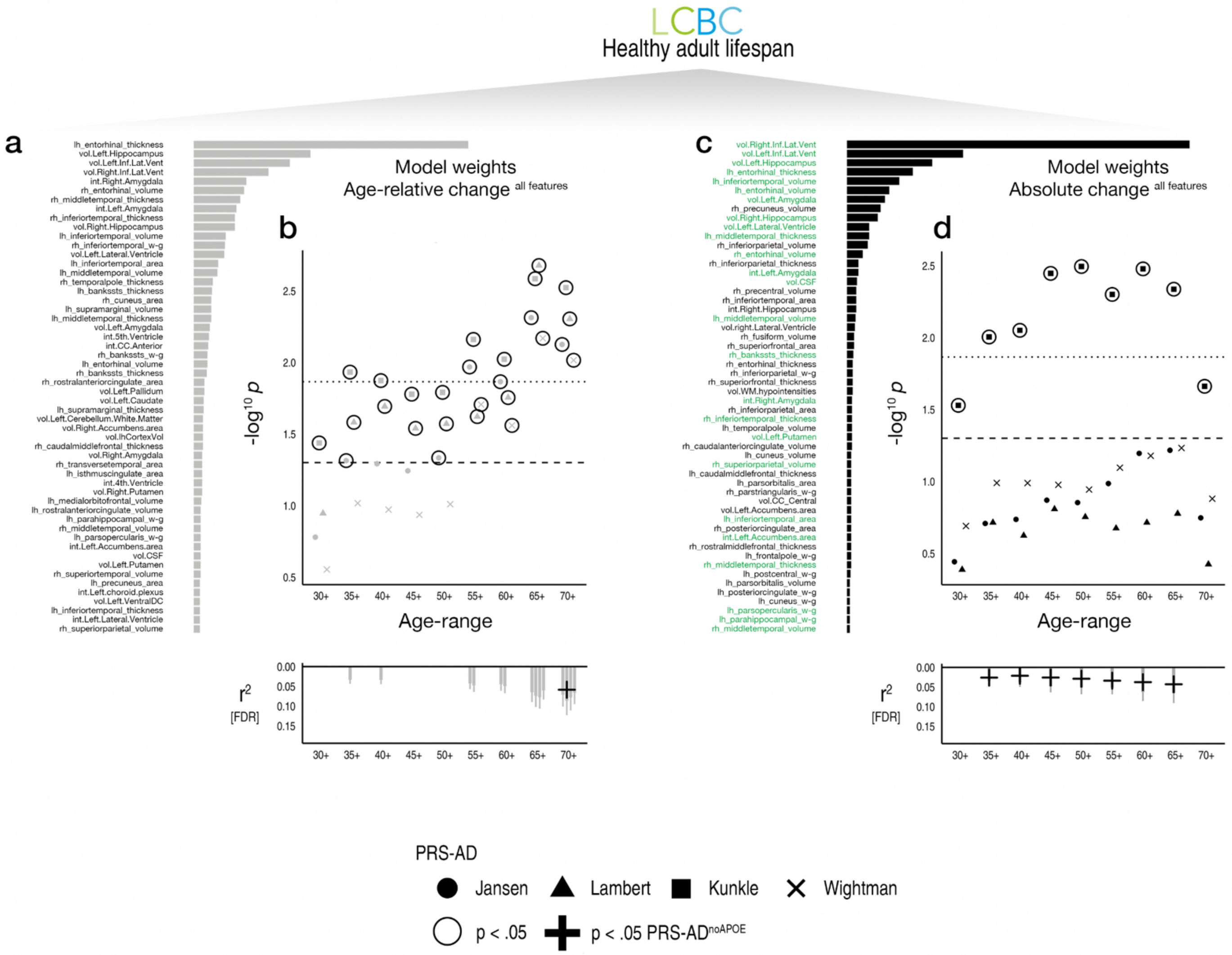

**SI FIGURE 5**

**ADNI-derived ML models applied to the healthy adult lifespan. a** Top features for classifying *AD-long* from *NC-long* individuals in ADNI data based on age-relative change (as in Fig. 4A). We directly applied the ADNI-derived model weights to the LCBC healthy adult lifespan dataset. This prediction therefore incorporates information from the weights of all 364 features. **b** PRS-AD associations in the LCBC healthy adult lifespan dataset after predicting with the learned model weights from the ADNI-derived model (i.e. LCBC as test data). The dependent variable is the model-implied log odds of having AD ($probAD^{relChange}$) based on age-relative change. Datapoints show (-log10) p-values for PRS-AD associations with $probAD^{relChange}$, tested at progressively older age-ranges, for all four scores. Dashed line indicates p=.05, and datapoints with black stroke denote significant PRS-AD associations at p<.05. Datapoints above the dotted line are significant at p(FDR)<.05 (FDR-correction applied across all 72 PRS-AD associations tests performed in this analysis). Bottom plot shows partial $r^2$ for PRS-AD where the association survived FDR-correction. Where FDR-corrected significant, we retested the association after removing APOE ($PRS\text{-}AD^{noAPOE}$). Partial $r^2$ of $PRS\text{-}AD^{noAPOE}$ is depicted by a black cross if the association remained significant ($p < .05$). **c** Top features for classifying *NC-long* from *AD-long* individuals in ADNI data based on absolute change values. Green text indicates whether the feature was present or not in the list of top features from the model shown in a. **d** PRS-AD associations in the LCBC dataset after predicting with ADNI-derived model weights based on absolute change. Here, the dependent variable is the model-implied log odds of having AD based on absolute change ($probAD^{absChange}$). Bottom plot shows partial $r^2$ for PRS-AD where the association survived FDR-correction, and partial $r^2$ of $PRS\text{-}AD^{noAPOE}$ is depicted by a black cross if the association remained significant after removing APOE ($p < .05$). Note that fewer associations were evident using absolute change, suggesting models based on age-relative change may be superior at capturing individual differences in brain ageing. Error bars depict 95% CI. lh=left hemisphere, rh=right hemisphere, vol=volume (subcortical); int=intensity (subcortical); w-g=grey/white matter contrast. Subcortical features (aseg) are delineated with ".", whereas cortical features (aparc) are delineated with "_".

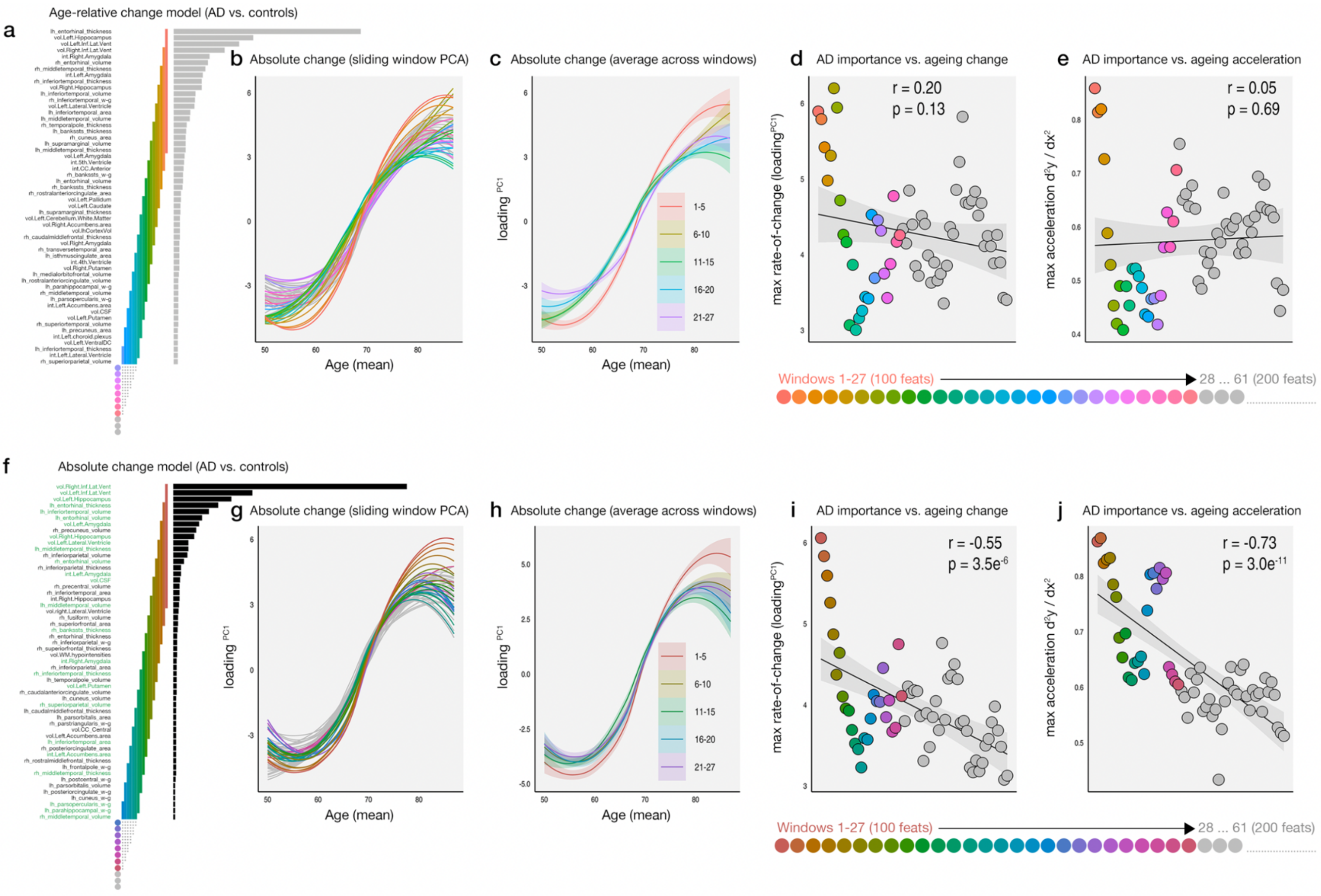

**SI FIGURE 6**

PCA-based sliding window trajectory analysis in the LCBC adult lifespan discovery sample. Within the 50-89 years age-range, we ran a sliding window PCA, iteratively calculating PC1 across 20 features with a step size of 3. We calculated the trajectory of absolute change within each window. Colours denote the selection of features across which we calculated PC1 of absolute change (here including hippocampal and amygdala volumes) and link with the subsequent plots. Feature selections and colours high up in the importance matrix thus represent those that contribute most to separating AD patients from controls (i.e. AD importance), whereas selections that are comparatively lower down contribute less to this prediction. **a** Top features for classifying *AD-long* from *NC-long* individuals in ADNI data based on age-relative change (as in Fig. 4A). **b** Absolute change as a function of mean age across timepoints within each window. Since the y-axis in b, c, g and h represents change, the slope of the curves represents acceleration. Feature selections across the top 100 features are shown in colour (27 windows), whereas trajectories in grey represent feature selections beyond the first 100 features (up to 200 features; 61 windows). Note that features most important for separating AD patients from controls showed the highest rate-of-change and steepest acceleration in healthy individuals. **c** As in b, except the trajectories are averaged across the feature windows denoted in the key to better show their differences. Ribbons depict 95% CI. **d** Maximum rate-of-change and **e** acceleration of the brain aging trajectories in healthy individuals, plotted against AD-importance (x-axis shows the feature selections and thus implicitly represents AD-importance). Note that the plots confirm that maximal rate-of-change and acceleration in healthy individuals is found in features that are most important for separating AD patients from controls. **f-j** As in a-e, except feature selections follow the order of importance of features for classifying *AD-long* from *NC-long* individuals based on absolute change instead (i.e. as in SI Fig. 5C). Again, maximal rate-of-change and acceleration in healthy individuals was found in features most important for separating AD patients from controls.

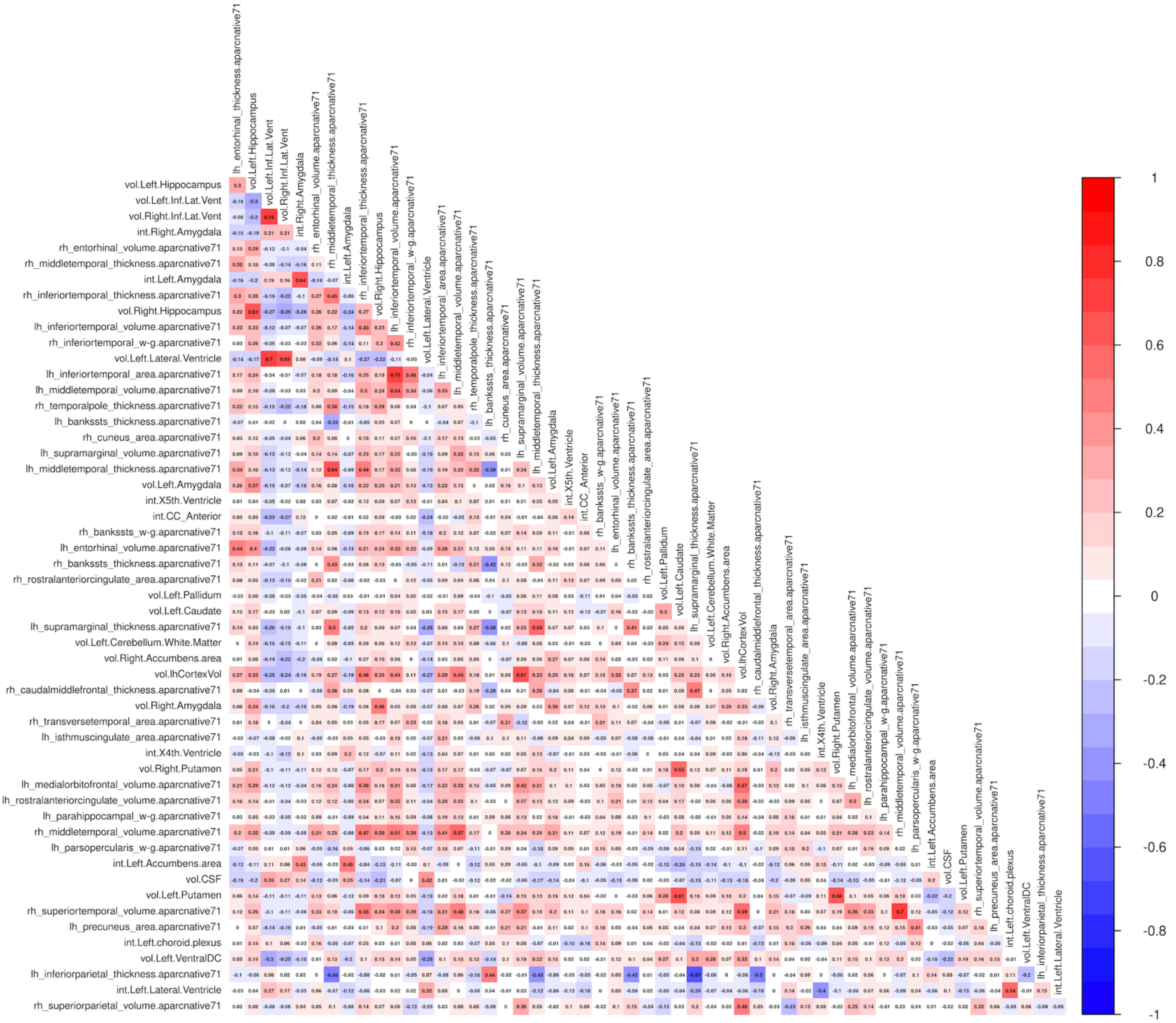

**SI FIGURE 7**

**Correlations between age-relative change estimates in the top 54 AD-accelerated features in LCBC healthy adult lifespan data**. Correlations are shown for the age-range 50-89 years. 54 features are shown due to the inclusion of hippocampal and amygdala volumes not otherwise included in PC1$^{relChange}$.

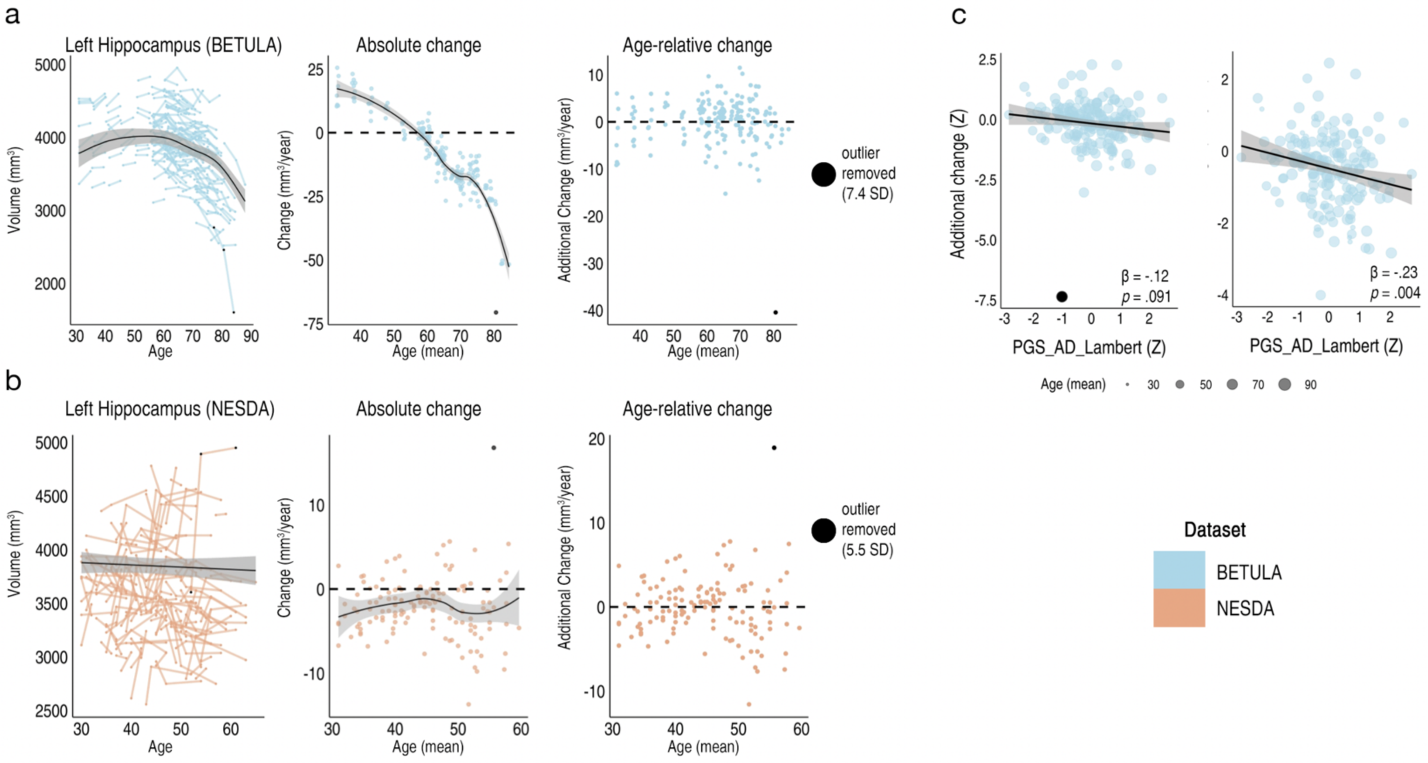

**SI FIGURE 8**
Initial GAMM trajectory analysis performed separately in the BETULA and NESDA longitudinal replication samples (max 3 timepoints) revealed one strong outlier in the left hippocampal slope data in each sample (datapoint in black indicates the large negative outlier in BETULA data [7.4 SD; top row], and the large positive outlier in NESDA data [5.5 SD; bottom row]). **a-b** These outliers are shown for each of their timepoints in the adult lifespan trajectories (leftmost plots), and absolute (middle plots) and age-relative change estimates (i.e. individual-specific slopes; rightmost plots) in each sample. **c** In BETULA, the strong outlier had a large influence on the tested PRS-AD association with age-relative change, here shown before (β = -.12, *p*=.091; left plot) and after removal of this outlier (β = -.23; p=.004; right plot; shown for one example score, though the data with all four tested scores was similar). The two samples were then collated into a single replication dataset and these outliers were removed.

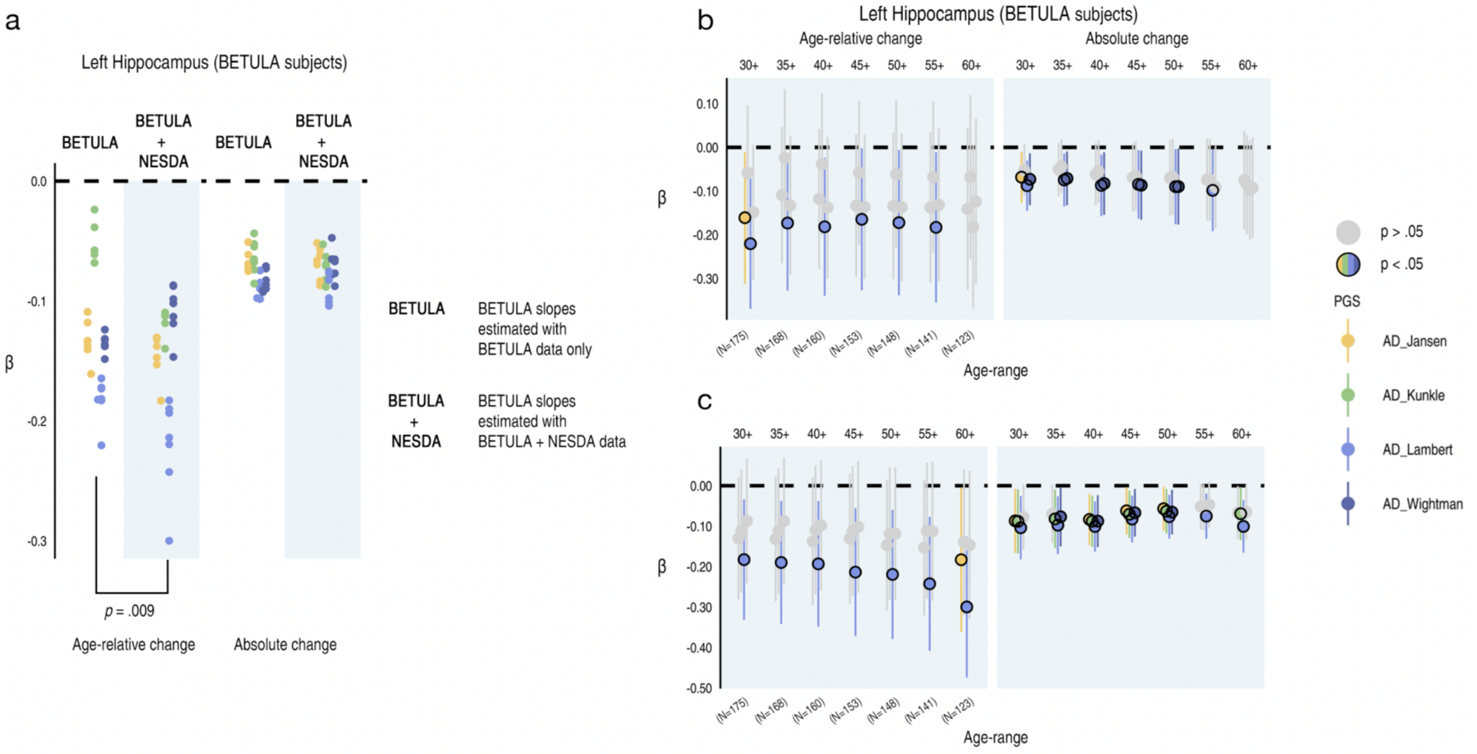

**SI FIGURE 9**

**a** In the same individuals in BETULA data (i.e. all BETULA subjects), PRS-AD beta estimates with age-relative change in left hippocampus were significantly lower (*p* = .009 [one-sided]) when their individual-specific slopes were estimated together with NESDA data, relative to when estimated using only BETULA data. **b** Beta estimates and significance indicator (black stroke denotes p<.05) for individual-specific slopes (BETULA subjects) estimated from a GAMM using only BETULA data. **c** Beta estimates and significance for individual-specific slopes (same subjects) estimated from a GAMM across BETULA and NESDA data. Though the number of significant associations with age-relative change was roughly the same, beta estimates were significantly lower (a), and more significant associations were evident with absolute change. Hence, including more longitudinal observations in the GAMM helped optimize the estimation of individual-specific slopes for all, boosting the power to detect PRS-AD associations in the same individuals. Error bars denote 95% CI.

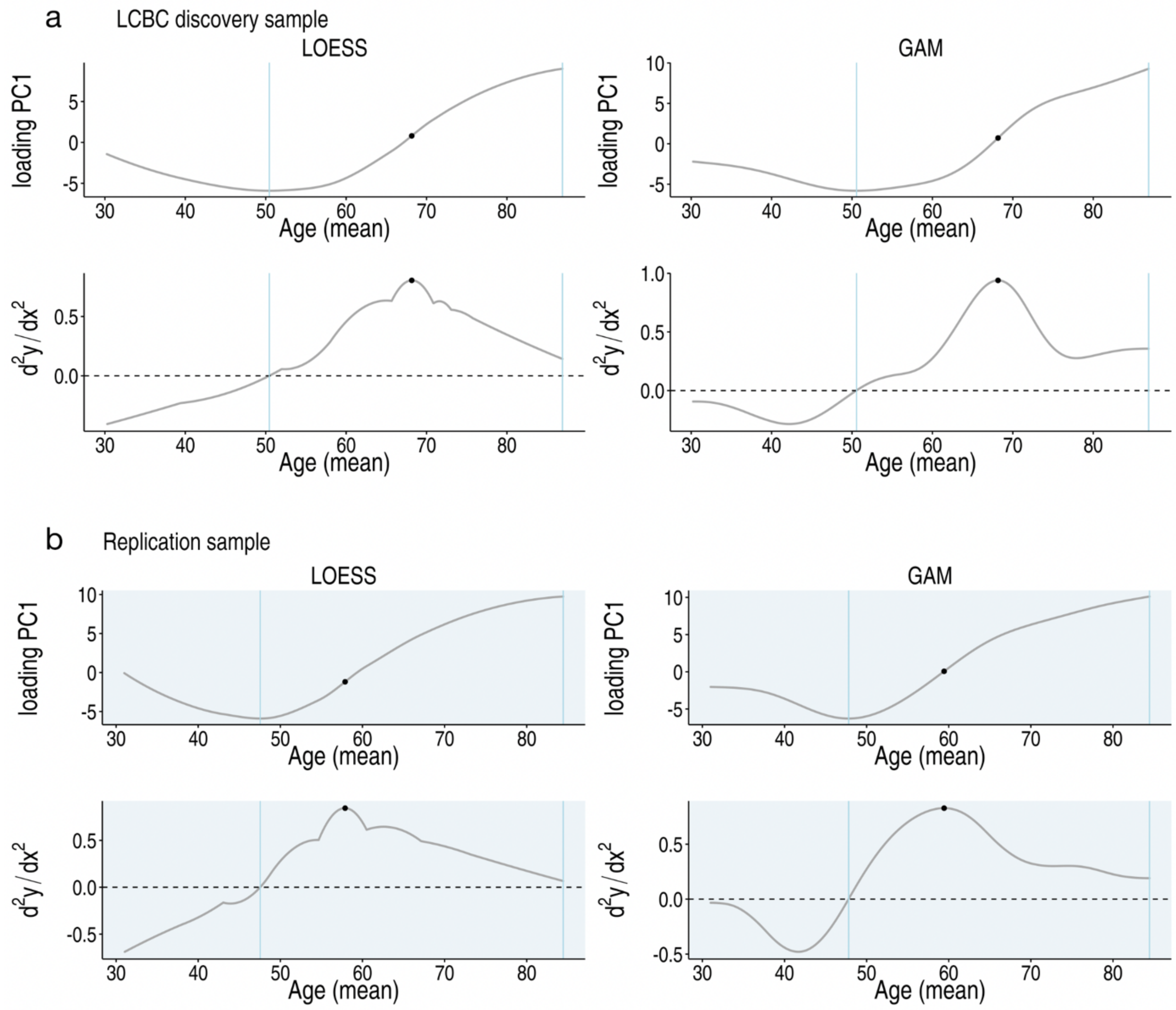

**SI FIGURE 10**

**Multivariate change in AD features (PC1$^{absChange}$) in healthy adults**

**a** For the LCBC adult lifespan discovery sample, top row shows the rate of absolute change (PC1$^{absChange}$) which reflects the first derivative (i.e., the y-axis represents rate-of-change; see Methods). Bottom row shows the derivative of this curve, which therefore represents acceleration (i.e., second derivative). Left column: estimated using a Locally Estimated Scatterplot Smoothing (LOESS) model (as in Fig. 4 and Fig. 5). Right column: estimated using a General Additive Model (GAM). The first and second blue line markers indicate the onset of negative change (crossing to positive on the second derivative) and the point of maximum rate-of-change in AD features in healthy adults. Black points indicate the point of maximum accelerated change. **b** As above, shown for the adult lifespan replication sample.

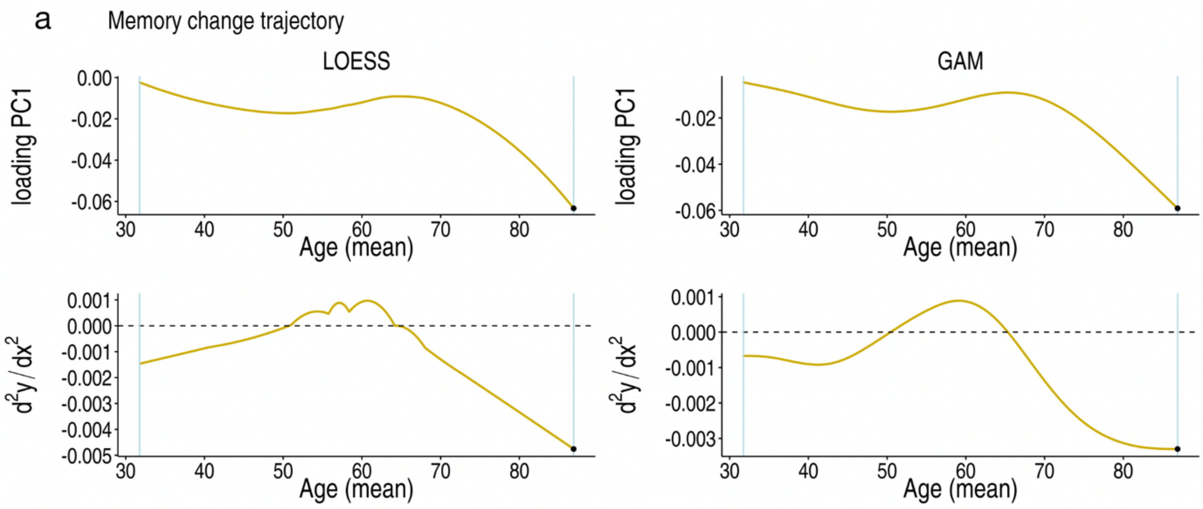

**SI FIGURE 11**

**Memory change trajectory across the healthy adult lifespan**
**a** Top row shows the rate of absolute memory change which reflects the first derivative (i.e., the y-axis represents rate of change; see Methods). Bottom row shows the derivative of this curve, which therefore represents acceleration (i.e., second derivative). Left column: estimated using a Locally Estimated Scatterplot Smoothing (LOESS) model (as in Fig. 4 and Fig. 5). Right column: estimated using a General Additive Model (GAM). The first and second blue line markers indicate the onset of negative memory change (estimated at the minimum age [mean] of our sample) and the point of maximum rate-of-change in memory in healthy adults (estimated at the maximum age [mean] of our sample). Black points indicate the point of maximum accelerated negative memory change.

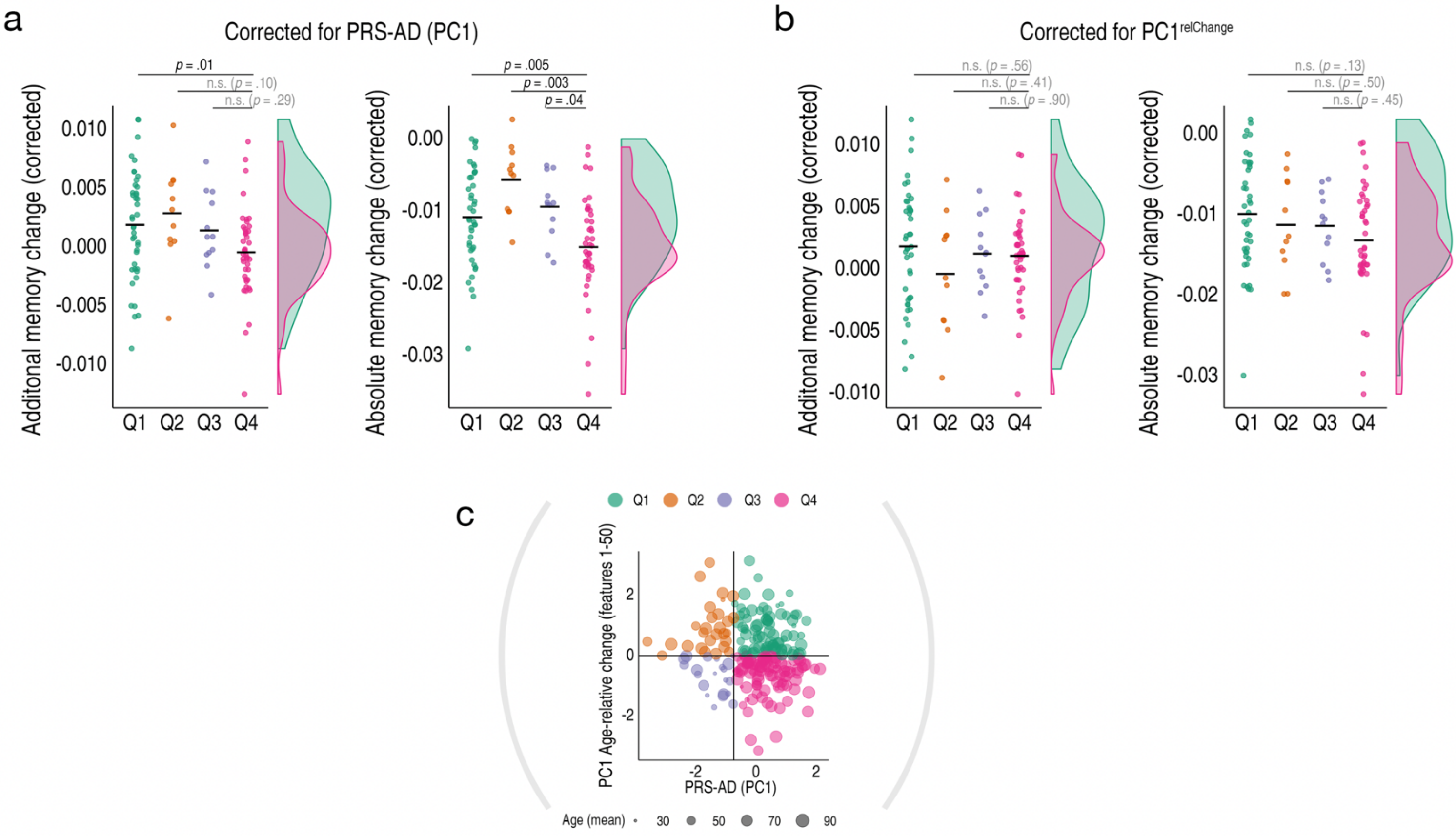

**SI FIGURE 12**

**Memory change results for alternative models correcting for additional covariates a.** Correcting for group-differences in genetic risk (PRS-AD; PC1 across four scores), and **b.** group-differences in multivariate brain change across AD-accelerated features ($PC1^{relChange}$). The reported group-differences in memory change persisted when additionally controlling for group-differences in genetic risk (similar results were found correcting for the number of APOE e4 alleles; see Results) but not when correcting for group-differences in brain change (other covariates: mean age, sex, number of timepoints, interval between first and last timepoint). **c.** Shown again for comparison, the association between the principal component across the four PRS-AD scores and the principal component of age-relative change across the first 50 ADNI-derived features (listed in Fig. 4A) that was used to define four quadrant-groups representing the conjunction of brain and genetic risk factors (as in Fig. 4D).

| | | | | N Timepoints | | | | | | | | | | | | | Mean MMSE |
|---|---|---|---|---|---|---|---|---|---|---|---|---|---|---|---|---|---|
| Dataset | Cohort | N Unique | N obs | 2 | 3 | 4 | 5 | 6 | 7 | Mean Time Interval (SD) | Interval Range | Mean Age (SD) | Age-Range | Sex (f/m) | N Genetic | Age < 60 (N obs = 405) | Age > 60 (N obs = 909) |
| Discovery | LCBC | 420 | 1430 | 135 | 147 | 45 | 26 | 60 | 7 | 2.1(2.8) | 0.14 – 11.1 | 63.7 (14.4) | 30.1 – 89.4 | 248 / 172 | 229 | 29.3 (0.8) | 28.8 (1.2) |
| Replication | BETULA | 182 | 449 | 97 | 85 | | | | | 3.0 (2.7) | 3.5 – 7.7 | 64.3 (11.9) | 30.9 – 87.9 | 85 / 97 | 175 | | |
| | NESDA | 138 | 331 | 83 | 55 | | | | | 2.5 (3.3) | 1 – 10 | 45.1 (7.9) | 30 – 65 | 91 / 47 | 118 | | |

**SI TABLE 4**
Description of the LCBC discovery healthy adult lifespan sample and the two Lifebrain cohorts that comprised the replication sample. Intervals are in years.

| Sample | Scanner | Tesla | Sequence parameters |
|---|---|---|---|
| LCBC | Avanto Siemens | 1.5 | TR: 2,400 ms, TE: 3.61 ms, TI: 1,000 ms, flip angle: 8°, slice thickness: 1.2 mm, FoV: 240 × 240 m, 160 slices, iPat = 2 |
| | Avanto Siemens | 1.5 | TR: 2,400 ms, TE = 3.79 ms, TI = 1,000 ms, flip angle = 8, slice thickness: 1.2 mm, FoV: 240 × 240 mm, 160 slices |
| | Skyra Siemens | 3.0 | TR: 2,300 ms, TE: 2.98 ms, TI: 850 ms, flip angle: 8°, slice thickness: 1 mm, FoV: 256 × 256 mm, 176 slices |
| BETULA | Discovery GE | 3.0 | TR: 8.19 ms, TE: 3.2 ms, TI: 450 ms, flip angle: 12°, slice thickness: 1 mm, FOV 250 × 250 mm, 180 slices |
| NESDA | Phillips | 3.0 | TR: 9 ms; TE: 3.5 ms; slice thickness: 1 mm, FOV: 256 x 256 mm, 170 slices. |

**SI TABLE 5**
**MRI parameters**
FoV = field of view, iPat = in-plane acceleration, TE = echo time, TI = inversion time, TR = repetition time.

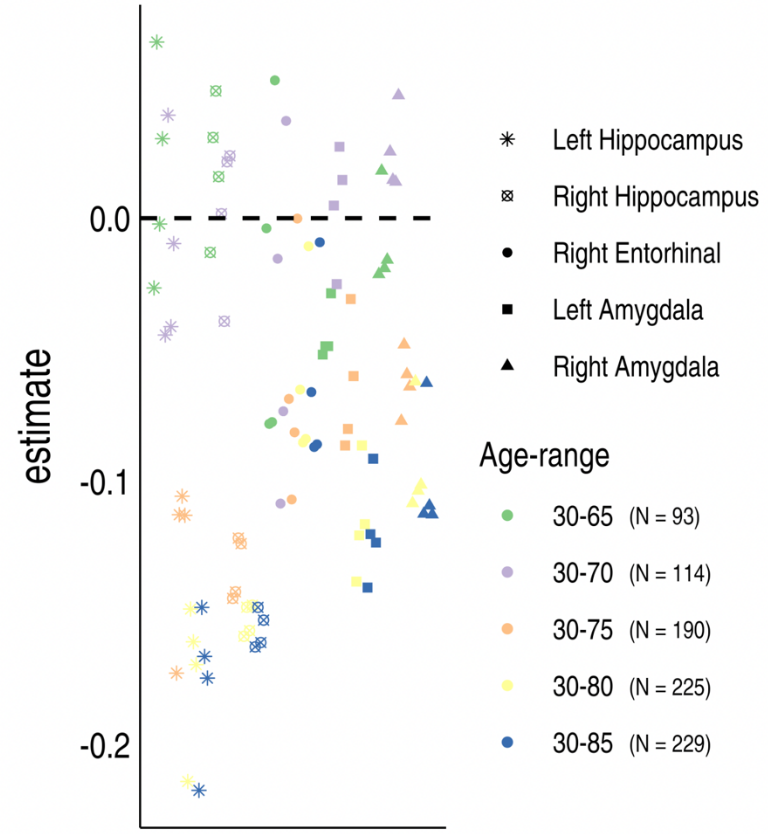

**SI FIGURE 13**
Beta estimates for PRS-AD associations with age-relative change in early Braak stage regions, after progressively disgarding the data from the oldest participants (age-range; shown for Braak regions wherein PRS-AD associated with change in the main analyses; LCBC discovery sample; covariates mean age (across timepoints), sex, N timepoints, interval between first and last timepoint, and 10 genetic PCs). The data indicated the associations were not driven by only the oldest-old, though older adults likely contributed more of the individual differences in brain change signal.

**SUPPLEMENTARY NOTE**

**LCBC Samples**
The main discovery sample consisted of magnetic resonance imaging (MRI) data collected across 5 projects at the Center for Lifespan Changes in Brain and Cognition (LCBC; Department of Psychology, University of Oslo). For the brain change analysis, to optimize individual-specific age-relative change estimates for all individuals, we used as many longitudinal observations as we could gather. Notably, the samples described below were all used in the brain change analysis, but many of these individuals had no genetic observations, and thus were not included in the PRS-AD association tests (see below). Brain change data over the following projects were used, and the number of individuals included in the genetic analysis [max N = 229] is also given per project. For the memory change analysis, to optimize individual-specific memory slopes for all we also used as many usable longitudinal memory observations as we could. However, we discarded all data for individuals involved in memory training projects with on-off designs over time, which likely to induce non-linear effects on the individual-specific memory slopes.

**Cognition and Plasticity Through the Lifespan (MemP):**
For the brain change analysis, **537** scans of 185 individuals (mean age = 59.5 ± 13.0, age-range = 30.6 – 89.4, females = 110, 2–7 timepoints [TP's]), collected on 3 different scanners were used (N 2TP's = 57, 3TP's = 93, 4TP's = 33, 5TP's = 1, 7TP's = 1). The number of scans used here collected on each scanner was 489 (1.5T Avanto), 46 (3.0T Skyra) and 2 (3.0T Prisma), respectively.
*For the genetic analysis, **81** individuals originated from this project.*
*To estimate individual-specific memory slopes, **522** observations from **185** individuals originated from this project,*

This is an ongoing longitudinal study where cognitively healthy adults have undergone MRI scanning and neuropsychological evaluation. The study consists of four main recruitment waves (though note that the number of longitudinal timepoints ranges from 2-7). New participants were recruited at waves 1, 3 and 4. Participants were scanned up to 11.1 years after the initial scan. The interval between waves was approximately 3.5, 4.4 and, 1.7 years, respectively. Data acquisition took place between 2006 and 2023 at the center for LCBC. For more details see [1,2]. Volunteers were initially recruited by newspaper advertisements and later contacted by mail for follow-ups. MRI sequences were acquired across three scanners (1.5T Avanto, 3.0T Skyra, 3.0T Prisma)

**Constructive Memory (MemC):**
For the brain change analysis, **85** scans of **42** individuals (mean age = 53.9 ± 13.4, age-range = 30.5 – 80.8, females = 24, 2-3 timepoints [TP's]), collected across two scanners were used (N 2TP's = 41, 3TP's = 1). The number of scans here collected on each scanner was 84 (3.0T Skyra) and 1 (3.0T Prisma), respectively.
*For the genetic analysis, **30** individuals originated from this project.*
*To estimate individual-specific memory slopes, **80** observations from **41** individuals originated from this project.*

This project is a cross-sectional study where cognitively healthy adults underwent an fMRI source-item memory task. The protocol also included MRI scanning and neuropsychological evaluation. The sample was collected at the center for LCBC. The project is nested within the longitudinal Cognition and Plasticity through the Lifespan project. Data acquisition took place between 2013 and 2015. MRI sequences were acquired with a 3.0T Skyra scanner. See [3,4] for more details.

**Neurocognitive Plasticity (NCP):**
For the brain change analysis, **637** scans of **130** individuals (mean age = 70.5 ± 12.3, age-range = 30.1 – 84.0, females = 79; 2-3 timepoints [TP's]), collected on a single scanner (3.0T Skyra) were used (N 2TP's = 17, 3TP's = 11, 4TP's = 12, 5TP's = 24, 6TP's = 60, 7TP's = 6).
*For the genetic analysis, **117** individuals originated from this project.*
*To estimate individual-specific memory slopes, **0** observations originated from this project (due to the complex on-off memory training design over time likely to induce non-linear effects upon individual-specific memory slopes; see below)*

This study consists of an experimental project of memory training with the "method of loci", employing an on-off training design over time. The study includes two groups of participants (young and old) that underwent an ABAB design where a batch in each group started with a resting condition, and the other started with memory training (after a baseline test). The study also includes an active group without memory training. The participants were initially scanned up to 6 times, five of them before/after a block of training while the sixth time point consisted of a follow-up ≈ 2 years after the intervention (note that several participants also have a seventh timepoint). Participants were recruited through newspaper and web page adverts and were screened with a health interview. Participants were required to be either young or older (in or around their 20s or 70s, respectively) healthy adults. Data acquisition took place between 2013 and 2018. MRI sequences were acquired with a 3.0T Skyra scanner. See [5,6] for more details.

**Method of Loci (Loci):**
For the brain change analysis, **112** scans of **41** individuals (mean age = 63.2 ± 9.1, age-range = 41.9 – 82.6, females = 22; 2-5 timepoints [TP's]), collected across two scanners were used (N 2TP's = 13, 3TP's = 27, 5TP's = 1). The number of scans used here collected on each scanner was 110 (1.5T Avanto) and 2 (3.0T Skyra), respectively.
*For the genetic analysis, **1** individual originated from this project (individual with 5 timepoints; see below).*
*To estimate individual-specific memory slopes, **111** observations from **41** individuals originated from this project (due to the simple pre-post memory training design [see below], individual-specific slopes are likely to be linear; note also that as only 1 individual here had genetic data, results of the longitudinal memory change analyses in the main paper – requiring individuals with both genetic and memory data – are unaffected by including this project, except insofar as the additional longitudinal observations may boost power to estimate individual-specific memory slopes for all; see below)*

This study consists of an eight-week simple pre-post memory training experiment focused on improving verbal recall memory by implementing the mnemonic technique "method of loci". Participants were scanned three times as part of the project: pre and post-training and a follow-up after 5 years. The number of timepoints included here therefore typically ranges from 2-3, though we note one participant originating from this project and included in the genetic analyses had 5 timepoints, as they were assimilated into (or later found to be a duplicate ID) of a later LCBC project (MemC). The full project included both healthy controls and memory clinic patients. Only data from healthy controls was used here. Healthy volunteers were recruited through a local newspaper ad, screened by a structured interview, and randomly assigned to either an intervention group or a control group serving as passive controls. Cognitive assessments and the memory training program were conducted at the center for LCBC. Data acquisition for waves 1 and 2 took place between 2007 and 2008 while the third wave was acquired in 2013. MRI sequences were acquired with a 1.5T Avanto scanner. See [7,8] for more details.

**Set to Change (S2C):**
For the brain change analysis, **59** scans of **22** individuals (mean age = 44.6 ± 16.8, age-range = 30.2 – 79.0, females = 13), collected on a single scanner (3.0T Prisma) were used (N 2TP's = 7, 3TP's = 15).
*For the genetic analysis, **0** individuals originated from this project (individuals were twins; see below).*
*To estimate individual-specific memory slopes, **0** observations originated from this project (due to the complex on-off memory training design over time likely to induce non-linear effects upon individual-specific memory slopes; see below)*

This is the first LCBC twin project, where twins were invited to participate, as recruited through either the Norwegian twin registry or via ads on social media platforms. The project consists of a targeted experimental approach to test differences in neurocognitive plasticity by training of younger and older adult mono- (MZ) and dizygotic (DZ) twins. The project employs a memory intervention using navigation training while cycling in virtual reality through a bespoke virtual city. Twins are assessed with brain MRI, and cognitive measures at multiple time points across 2.5 years pre- and post-a 10 week intervention in a AB/BA crossover design. Data acquisition began in 2019 and is ongoing. MRI sequences were acquired with a 3.0T Prisma scanner. Of the 22 individuals included in the brain change analysis here, 14 were MZ whereas 8 were DZ twins.

**Common information:**
Common exclusion criteria across all LCBC projects are outlined in the main paper. Written informed consent was obtained from all participants, and all studies were approved by the Regional Ethical Committee of South Norway.
